# Joint dog and wolf genealogies reveal the evolution of the canine genome

**DOI:** 10.64898/2026.06.18.729863

**Authors:** Jasmin Rees, Mattias Sherman, Daniel Teofilov, Aina Colomer i Vilaplana, Simon R. Myers, Anders Bergström, Leo Speidel

## Abstract

Dogs and their closest extant relative, the grey wolf, diverged around 30k years ago, but have since experienced complex histories of gene flow involving other canids, adaptive pressures due to close association with humans, and changing climates. We infer joint genealogies of dogs, grey wolves, and a coyote using available whole genomes to reconstruct the evolutionary forces shaping the dog genome. These genealogies reveal multiple strong mutation-rate pulses unique to dogs, including signals detectable across ancient dogs from the past 10,000 years. We further detect pervasive genealogical signatures of purifying selection and find that GC-biased gene conversion is a major driver of diversity patterns around gene promoters in dogs. We introduce a new genealogy-based selection scan, TwigScan, that computes time-stratified differentiation, increasing power over traditional F_ST_-based approaches. Applying this framework, alongside a second single-population test for detecting more recent selection within dogs, we identify multiple known and novel loci with signatures of positive selection. Among these, the region surrounding the amylase 2B locus shows evidence of introgression from a deeply divergent, unsampled canid lineage with divergence comparable to that of dholes. *AMY2B* duplications appear to occur exclusively on this introgressed haplotype which increased in frequency approximately 7,000–8,000 years ago, coinciding with increased reliance on starch-rich diets in human populations. Together, these results show how mutation-rate variation, gene conversion, selection, and inter-species gene flow have jointly shaped the dog genome, highlighting the power of genealogical approaches for resolving complex evolutionary histories.

## Introduction

The domestication of the dog represents one of the earliest known evolutionary transitions in response to close association with humans. Yet, many aspects of the full domestication trajectory, and associated genomic signatures, are still to be characterised. The grey wolf is the dog’s closest extant relative and is among the few large carnivorous mammal species that have maintained a wide geographic range despite extreme environmental changes, surviving past the Last Glacial Maximum (∼25kya). However, there remains no clear consensus on the geographic origins or timing of dog domestication, despite recent progress (Shannon et al. 2015; G.-D. Wang et al. 2016; Savolainen et al. 2002; Bergström et al. 2020; Gojobori et al. 2024; Bergström et al. 2022; Frantz et al. 2016; M. Zhang et al. 2025). Additional key questions include the extent to which genetic adaptations have enabled dogs to respond to changing environmental pressures, particularly introduced by living in close contact with humans. Hybridisation events amongst dogs and wolves, as well as with more distant canid relatives such as coyotes and jackals, may have facilitated such adaptation, but ultimately complicate canid phylogenetics (Gopalakrishnan et al. 2018; Lin et al. 2025). As such, canids are a compelling and challenging system for studying how demography, selection and introgression interact to shape genomes over a range of evolutionary timescales.

Dogs exhibit extraordinary phenotypic diversity despite their relatively low genetic diversity, with studies identifying artificial selection acting on paedomorphic traits and on genes related to body size, ear morphology and brain development within individual breeds (Meadows et al. 2023; Plassais et al. 2019; Cagan and Blass 2016; Pendleton et al. 2018; Pilot et al. 2021; Freedman et al. 2016). Other targets of selection in dogs also overlap with loci implicated in humans. Previous studies based on identifying loci of excess differentiation between dogs and wolves have detected signatures of adaptation at the pancreatic amylase *AMY2B* locus, which is involved in starch breakdown and may have been adaptive in the context of human-associated starch-rich diets (Axelsson et al. 2013; Maja Arendt et al. 2014). Humans also show a potentially adaptive wide range of copy number variation at the salivary amylase *AMY1* genes (Perry et al. 2007; Yilmaz et al. 2024; Bolognini et al. 2024). Similarly, the *EPAS1* locus is implicated in high-altitude adaptation in both human Tibetan populations, introgressed from Denisovans (Huerta-Sánchez et al. 2014), and Tibetan dogs and wolves, introgressed from a deeply divergent canid lineage (Li et al. 2014; M.-S. Wang et al. 2020; Miao et al. 2017).

Here, we reconstruct joint genealogies of dogs, grey wolves, and a coyote using two major genetic variation resources, the NHGRI Dog Genome Project (Plassais et al. 2019) and Dog10K project (Meadows et al. 2023). These genealogies enable us to disentangle the evolutionary forces shaping the canine genome, including demographic history, varying mutational processes, and natural selection, operating across the canine genome and along individual lineages through time. We identify several strong dog-specific mutation rate pulses, including a ubiquitous signature across dogs in the last 10k years and a separate strong signature unique to African dogs. We observe signatures of purifying selection around specific genes and widespread effects of GC-biased gene conversion impacting gene promoter sites, likely caused by a defunct *PRDM9* in canids (Axelsson et al. 2012).

To boost the power of selection scans previously used in dog studies, we introduce TwigScan, a cross-population selection scan that leverages inferred genealogies to calculate F_ST_ and improve power over traditional F_ST_-based approaches. Combined with a second single-population selection test, we identify eleven loci with evidence of positive selection, including cases of adaptive introgression, and more recent selection in domestic dogs. This includes strong evidence of positive selection around the *AMY2B* locus. We show that the selected haplotype was introgressed from an unknown ghost canid and *AMY2B* copy number expansions appear only on this introgressed haplotype, likely conferring an adaptive advantage related to starch digestion in post-Neolithic human-associated environments.

We show support for our genealogical inferences across two different independent datasets aligned to different reference genomes, complemented by further independent analyses that do not directly rely on genealogies and integrated with insights from ancient DNA. Our analyses here demonstrate the power of genealogical approaches in reconstructing complex evolutionary histories in a challenging system and reveal the confluence of evolutionary pressures acting on the canine genome.

## Results

### A joint genealogy of dogs, wolves, and coyotes

We inferred genealogies using the software Relate (Speidel et al. 2019) for two of the largest currently available genetic variation datasets of dogs and wolves, the NHGRI Dog Genome Project comprising 722 whole genomes (abbreviated 722g) (Plassais et al. 2019) and the Dog10K consortium data (Dog10K) (Meadows et al. 2023; Ostrander et al. 2019). For the 722g dataset, we downsampled breed dogs and co-analysed these with all available grey wolf genomes and one coyote (SI Table 1). For the Dog10K dataset, we selected 281 village dogs and 57 grey wolf genomes (SI Table 2). We therefore retained 135 and 338 non-overlapping samples from each dataset (Supplementary Information, SI Figure 1, 2), aligned to two different reference genomes, canFam3.1 and canFam4.

**Figure 1.**
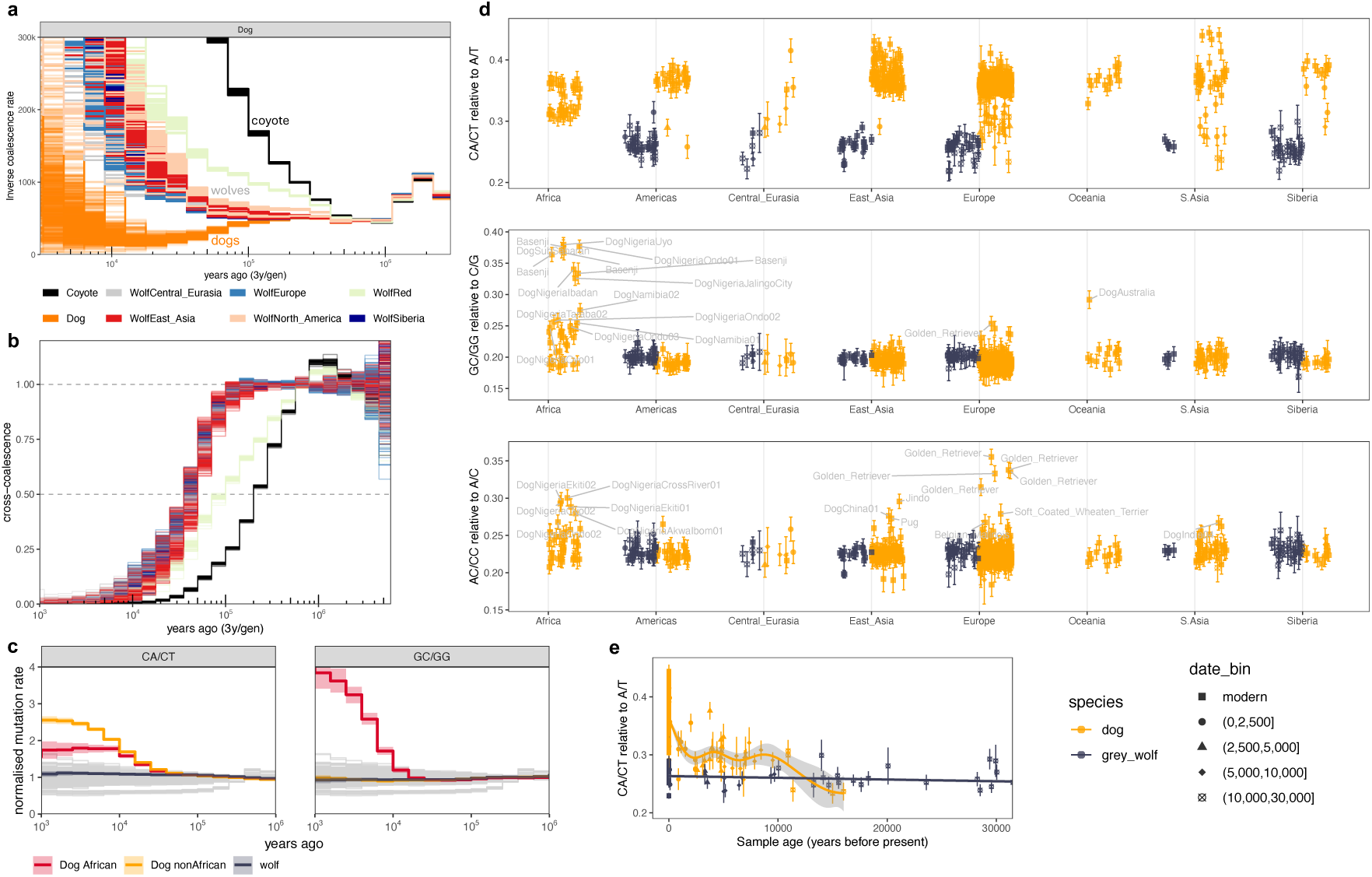
Canine demography and mutation rate pulses. **a**, Inverse coalescences rates between dogs and other canids, grouped by breed or region (1 coyote, 1 red wolf, 19 grey wolf, and 41 dog groups). **b,** Relative cross coalescence rates between dogs and wolves or a coyote, coloured by super-population assignment in a. **c,** Relate-inferred mutation rates of sequence contexts CA/CT and GC/GG, relative to the average mutation rate. Colours differentiate mutation rates in African dogs, non-African dogs, and grey wolves. Grey lines show trajectories for all remaining trinucleotide mutation rate contexts for each of the three species. All values in this figure were computed on the genealogies of the 722g data set and SI Figure 3 shows all 96 trinucleotide categories for the Dog10K and 722g datasets. **d,** Mutation rate pulse quantified using genotypes carried by each modern and ancient sample. We computed the ratio of mutations in context CA/CT, GC/GG, or AC/CC relative to A/T, C/G, and A/C respectively, restricting to mutations younger than 30,000 years (Supplementary Information). For GC/GG and AC/CC we highlight all African dogs, and other dogs in the top 2%. We show two standard errors computed using a chromosomal block jackknife. **e,** Temporal trajectory of mutations CA/CT relative to all A/T quantified in ancient dogs and grey wolves.

**Figure 2.**
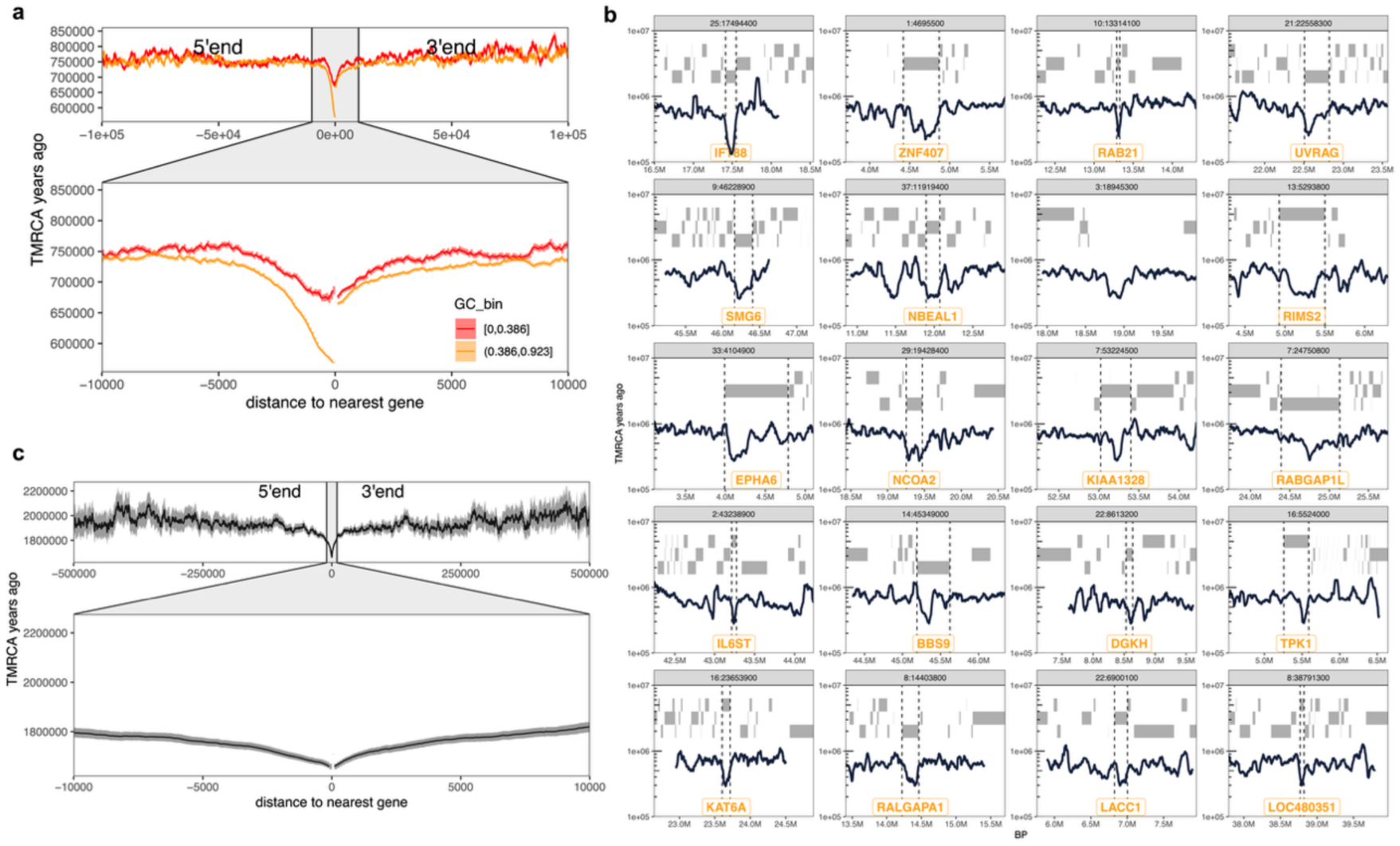
TMRCAs shaped by purifying selection and GC-biased gene conversion. **a**, TMRCA in dogs and wolves averaged by distance to the nearest gene and polarised by 5’ or 3’ end. We further stratify by GC content in each position bin. **b,** Top 20 youngest TMRCAs in dogs and wolves, of which 19 overlap a gene (**Methods**). Grey bars indicate protein coding genes and the focal genes are highlighted in orange. We use the 722g genealogies in a and b, and show results for Dog10K in SI Figure 7. **c,** Average TMRCA of humans in Relate genealogies reconstructed for populations CHB, GBR, and YRI of the 1000 Genomes Project. We compute distance to the 5’ or 3’ end of the gene and any bases falling inside a gene have distance zero. Error bars show 1.96 standard errors of the mean.

The inferred genealogies indicate an approximate split time of 10,000 generations (∼30,000 years) between dogs and grey wolves consistent with the literature (Skoglund et al. 2015; Bergström et al. 2022) and a ten-fold older split time with coyotes (**Figure 1a,b**). Cross coalescence rates are approximately symmetric between individual dog and wolf populations revealing no clear candidate wolf source population among currently available modern wolves. Previous split time estimates based on simulating histories using an Approximate Bayesian Computation approach have dated the coyote-wolf split time to 800,000-1M years ago using North American wolves and while accounting for gene flow between different coyote and grey wolf populations (Vilaça et al. 2023). We date the coyote-wolf divergence to around ∼200,000 years ago, although cross-coalescence rates start to diverge >1M years ago (**Figure 1b**). Our estimate may be consistent with a soft split with subsequent gene flow between grey wolves and the western coyote population from which our sample originates.

### Dog-specific mutation rate pulses

Germline mutations arise from a confluence of mutagenic processes that are influenced by sequence context, methylation status, parental age, and other factors (Seplyarskiy and Sunyaev 2021). The sequence context-dependent mutation spectrum has been shown to evolve across species (Beichman et al. 2023), however the underlying mechanisms driving variation in mutagenesis remain poorly understood.

We identify two strong sequence context dependent mutation rate pulses specific to dogs. By directly inferring nucleotide context-dependent mutation rates given genealogies (Supplementary Information, (Speidel et al. 2019)), we find a 2.5-fold increase in the CA to CT mutation rate and ACG to AAG in all modern dogs quantified relative to the long-term mutation rate of the respective single-nucleotide mutation category (**Methods**, **Figure 1c**, SI Figure 3). The pulse is not observed in grey wolves and the temporal mutation rate trend shows that this pulse occurred since the two species diverged (**Figure 1c**). To verify this pulse without relying on the full genealogical information, we additionally quantified this pulse by computing the proportion of CA/CT mutations relative to A/T mutations, restricting to mutations dated to within the last 30,000 years in the 722g genealogy (**Methods**, **Figure 1d**). This approach allowed us to use low coverage ancient dogs and wolves that were not incorporated into genealogies. The pulse appears significantly above the grey-wolf baseline in present-day dog populations and is also present in ancient dogs, although it appears weaker, likely in part attributable to the fact we do not call de-novo mutations in ancient individuals. Plotted as a time series (**Figure 1e**), the pulse is present across samples in the past 10k years, with two ∼10k-year-old dogs from northwestern Russia (Veretye) (Bergström et al. 2020; Feuerborn et al. 2021) being the oldest dog samples with a detectable signal. The signal is lowest and consistent with the grey wolf baseline in the three oldest dog genomes available, two >10k year old dogs from Turkey and a 15k year old dog from England (Bergström et al. 2026; Marsh et al. 2026), suggesting either regional stratification of the signal at that time or that the pulse had not yet started. Some >10k-year-old Siberian wolves appear to show weak evidence of the mutation pulse, however we caution on these results due to poor data quality. Overall, our results suggest that this signature is old, potentially starting in the early domestication period, but persisted and increased in strength over time.

**Figure 3.**
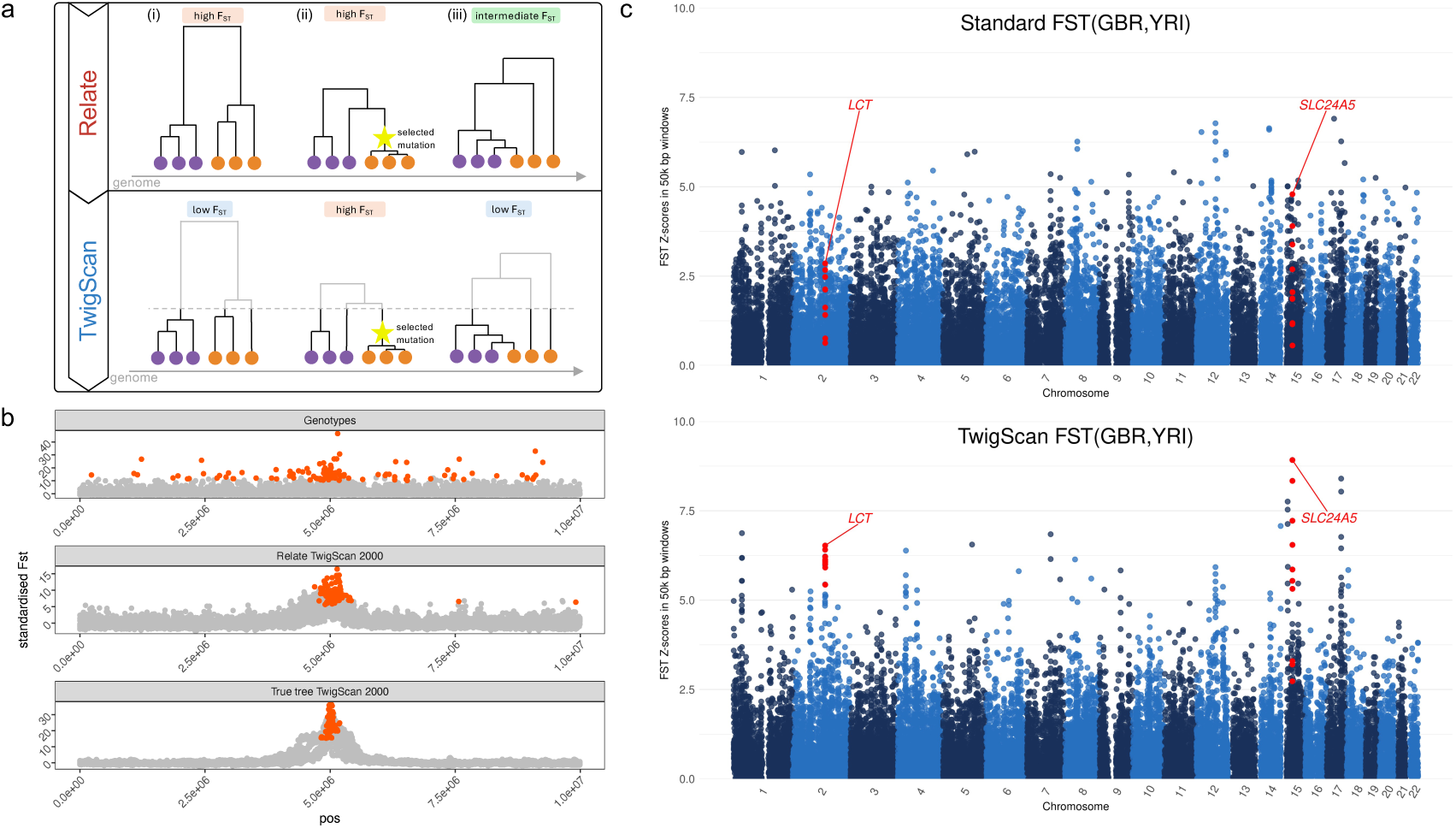
TwigScan performance. **a**, Schematic illustrating three example genealogies containing longer population-specific branches producing high F_ST_: (i) a long population-specific branch e.g. due to deep structure, (ii) recent positive selection producing a long population-specific branch, (iii) random coalescences producing imbalanced descendant make-ups, and the coalescent model generates longer branches at the top of the tree. TwigScan reduces F_ST_ in (i), (iii) but not (ii). **b**, TwigScan simulation (Supplementary Information) with a selected mutation in the middle of the genomic segment in one of two simulated populations. We set the selection coefficient to 1%. We compute F_ST_ between the two populations in genomic windows of 10kb, directly on genotypes and on Relate or true genealogies with a cutoff of 2000 generations. We show 10 replicates. In each replicate, we highlight windows with values in the top 1% in orange. **c**, TwigScan applied to 1000 Genomes Project human genetic variation data. We compute F_ST_(GBR, YRI) on genotypes or on Relate inferred genealogies in 50kbp windows with a cutoff 2000 generations ago (∼56ky).

Using both the direct genealogy estimate and mutation ratio approach, we also identify a remarkable ∼5-fold increase in the mutation rate of GC to GG and ACA to AGA almost entirely unique to African dogs (**Figure 1c, d**). Notably, this signature is observed strongest in the Basenji breed and village dogs from Nigeria and Namibia in the 722g dataset, while it appears strongest in village dogs from Kenya and Liberia in the Dog10K dataset, but interestingly is weaker in village dogs from the Congo, where Basenji dogs originate. We observe one additional case in an Australian dog with no apparent recent African ancestry and a weak signature in a Golden Retriever (**Figure 1c,d**). This signature appears in no currently available ancient dog genome, although we note that our dataset includes no ancient African dogs. Interestingly, we are unable to detect this signature in a low coverage 7.2k year old ancient dog from Israel (THRZ02), who appears ancestral to African dogs in previous analyses (Bergström et al. 2020). Finally, a subset of dogs, mostly Golden Retrievers and some African dogs also exhibit a slightly increased AC/CC mutation rate; this signature appears somewhat correlated to the GC/GG signature.

Both signatures are observed in both the 722g and Dog10K datasets, which contain independent samples and are aligned to separate reference genomes (SI Figure 3). In both datasets, we observe no notable mutation rate pulses in grey wolves. We further confirm that both signatures are observed at equal strength also for the reverse complement (e.g. CA/CT and TG/AG) and are therefore strand symmetric (SI Figure 3). In addition, the patterns persist when restricting to mutations carried by at least two dog or wolf samples, ruling out artefacts caused by singletons (SI Figure 4). Together, these observations make it highly unlikely that the signatures arise from sequencing artefacts.

**Figure 4.**
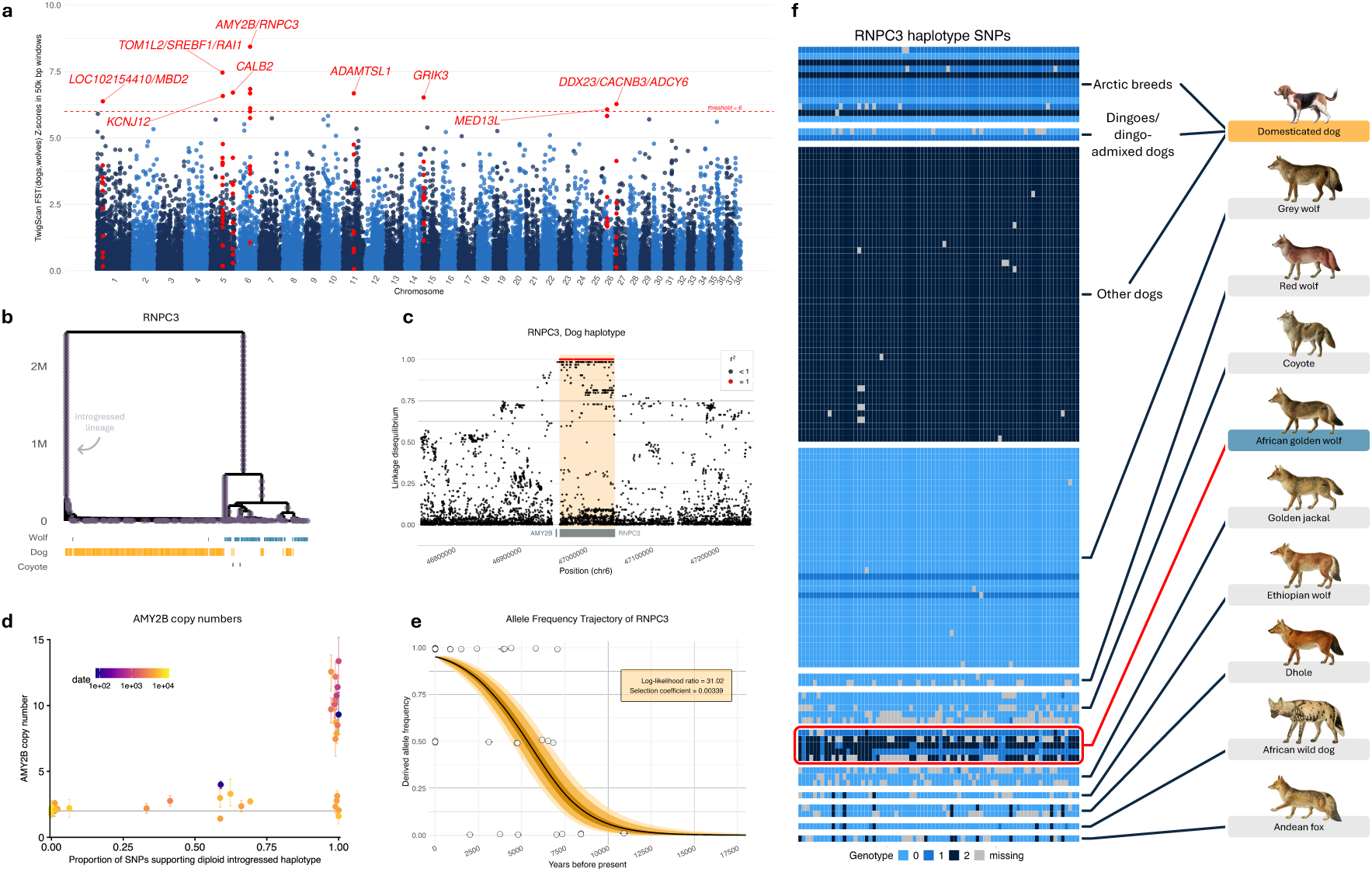
TwigScan applied to 722g dogs. **a**, Manhattan plot showing TwigScan F_ST_-scans (t = 14k gens) between dogs and grey wolves in the 722g dataset. Outlier regions are labelled with all protein-coding genes located in windows with z-scores above 6, and windows within 250kbp of the peak are highlighted in red. **b,** Relate genealogy at the *RNPC3* gene (chr6:47Mbp), corresponding to the highest F_ST_ peak in **a**. **c,** Introgressed haplotype identified by plotting SNPs in perfect LD, differentiating dogs from grey wolves around the *RNPC3* gene (Supplementary Information). **d,** *AMY2B* copy numbers plotted against introgression status in ancient dogs (**Methods**). Colour shows ancient DNA sample age. **e,** Allele frequency through time of the *AMY2B*/*RNPC3* haplotype inferred from Western Eurasian modern and ancient dogs using CLUES2 (Vaughn and Nielsen 2024) (Supplementary Table 6). **f,** Genotype matrix for the *RNPC3* gene. Only SNPs in perfect LD in modern dogs and grey wolves are shown, corresponding to mutations mapping to the top two lineages in **b.**

We observe no detectable localisation along the genome, and no effect with regard to distance to exons, with one notable exception. The GCG to GGG signature increases significantly with distance from exons (SI Figure 3). Importantly, these effects are not driven by GC-biased gene conversion (Galtier and Duret 2007), which we observe separately in the form of mutation rates towards G or C increasing and mutation rates towards A or T decreasing further in the past (SI Figure 5). This effect is driven by mutations towards G or C spreading more quickly in the population and therefore appearing more frequent and older, whereas mutations towards A or T decrease in frequency and are depleted in the more ancient past.

**Figure 5.**
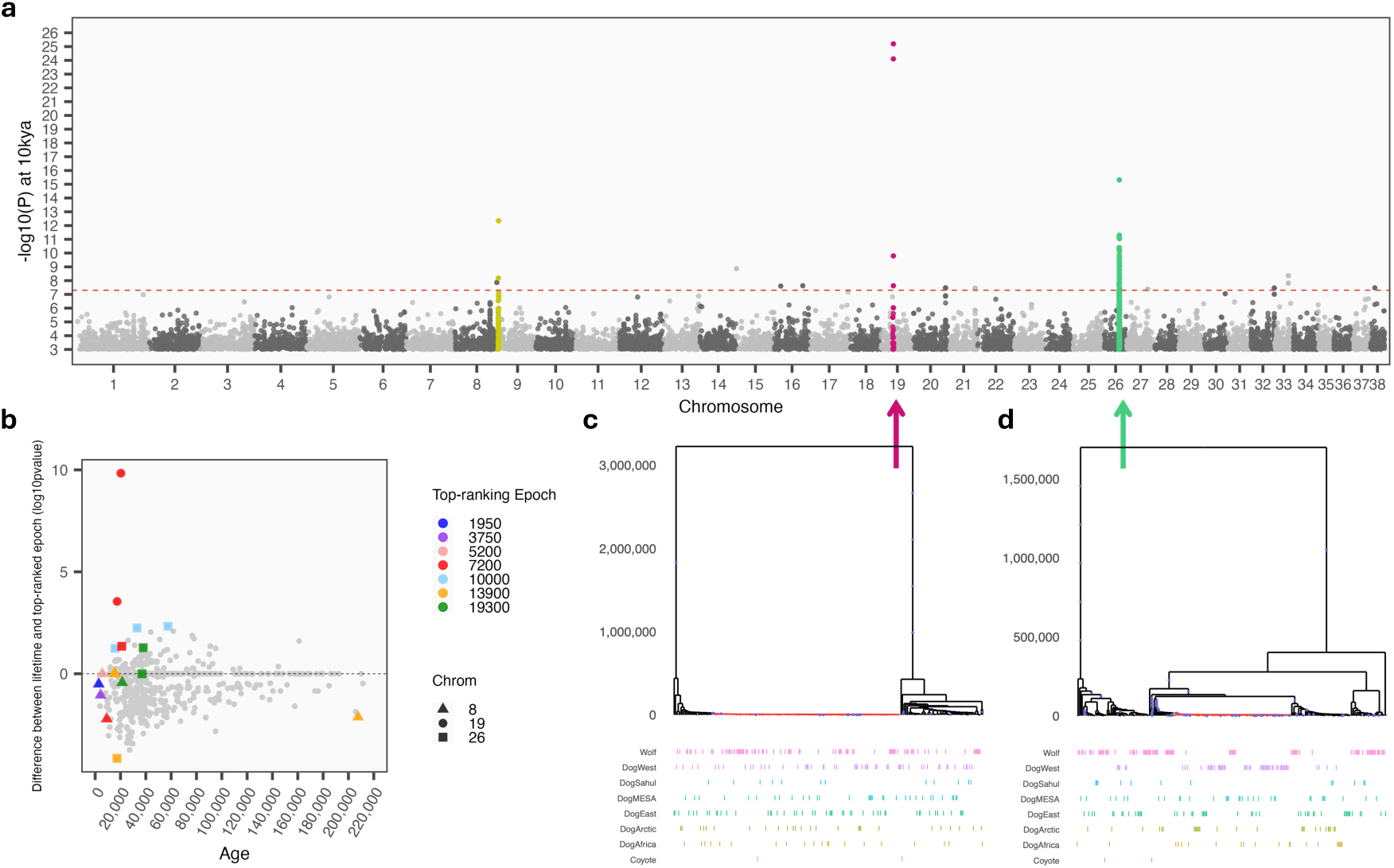
Relate selection test in 722g dogs. **a**, Manhattan plot showing the evidence of positive selection calculated using the Relate selection test over epochs, where we condition on 10kya. We highlight peaks on chromosomes 8, 19 and 26. **b,** Difference between the standard RST log₁₀(p) (testing lifetime selection) and the RST-over-epochs log₁₀(p) (testing recent selection), where we select the epoch with smallest p-value, plotted against inferred mutation age. The RST-over-epoch log₁₀(p) increases by as much as 10 on chr19. Mutation age is inferred from the 722g genealogies, where we used the midpoint of the branch. SNPs in the peaks of chr 8, 19, and 26 (indicated by shape; lifetime -log₁₀(p) > 7) are coloured by their respective epoch with strongest evidence of positive selection. **c,** Relate-inferred genealogy for chr19:20,248,039, **d,** Relate-inferred genealogy for chr26:26,293,281.

Examination of de-novo mutation calls in a pedigree study comprising 390 trios (S.-J. Zhang et al. 2025) showed that the mutation categories we identified were not enriched in excess de-novo mutations (SI Figure 6, Supplementary Information). We are likely reasonably well-powered to observe the dog-wide mutation signature CA to CT, which therefore suggests that this pulse is no longer active in dogs. For the African dog specific signature we rely on only a subset of de-novo mutations occurring in six Basenji trios and are likely underpowered.

To the best of our knowledge, this is only the second clear example of a temporally localised, context-specific germline mutation rate pulse in mammals. In humans, a different nucleotide context (TCC to TTC) was previously reported to have experienced a strong ∼2-3 fold mutation rate pulse 10k - 30k years ago, impacting primarily populations from Europe and South Asia, with potential origins in ancestors of early farmers (K. Harris 2015; Kelley Harris and Pritchard 2017; Speidel et al. 2021). Although numerous mutation signatures have been identified in the cancer literature, we have so far been unable to link the observed germline mutation pulses in humans and dogs to these known signatures (Alexandrov et al. 2020). One exception comes from laboratory mouse strains, in which a mutator allele has been identified linked to SBS18 (Sasani et al. 2022). Our findings suggest that germline mutation rates can be highly dynamic, but currently the underlying causes, and reasons for sequence context dependency remain unresolved.

### Species-wide sweeps and purifying selection

Rapid species-wide selective sweeps detected in population genetic variation data have typically remained elusive in mammalian species, including in humans (Mallick et al. 2016). An exception is that of grey wolves, where a previous study identified a strong sweep occurring 40k - 50k years ago by leveraging a 100,000 year ancient DNA time series (Bergström et al. 2022). The sweep region overlaps almost exactly with the *IFT88* gene, which is believed to impact cranio-facial morphology, and is fixed in present-day dog and wolf populations. Consistent with strong positive selection acting on this locus, the region was also inferred to have the youngest time to the most recent common ancestor (TMRCA) across grey wolves and dogs in Relate genealogies (Bergström et al. 2022).

To investigate if we can identify other loci with young TMRCAs, and therefore putatively under selection across grey wolves and dogs, we performed a genome-wide scan ranking the joint TMRCA of dogs and grey wolves along the genome (**Methods**). Encouragingly the youngest TMRCA corresponded to the *IFT88* gene in both the 722g and Dog10K datasets, aligning with previous evidence. To investigate other regions with young TMRCAs, we selected the top 20 youngest TMRCAs in the 722g genealogies.

Among these, 19 nearly perfectly overlapped with a gene and we observe a high correlation between the 722g and Dog10K datasets (**Figure 2b**, SI Figure 7). When compared to 1000 random regions passing genome mask filters, this overlap with genes was significant (Fisher’s exact test p-value = 2.5e-06). Among these 19 genes, seven (*ZNF407, UVRAG, RIMS2, NCOA2, IL6ST, KAT6A, RALGAPA1*) have a probability of loss-of-function intolerance score (pLI) of 1 in humans (gnomadv4.1; (Chen et al. 2024)), also constituting a significant enrichment (Fisher’s exact test p-value = 0.01703). This indicates that the low TMRCA of these loci are not exclusively driven by strong positive selection, as is the case for *IFT88*, but may be a result of strong purifying selection acting in highly constrained regions during canid evolution.

Averaged across the genome, we further observe a significant dip in TMRCAs around genes (**Figure 2a**). This is also observed in human genealogies (**Figure 2c**) which has been shown to be driven by background selection at linked loci (Hernandez et al. 2011). Contrary to humans however, the reduction in canid TMRCAs is asymmetric, with a much larger reduction observed at the 5-prime end of a gene in canids. This corresponds to an approximately 20% decrease in inferred TMRCAs compared to the genome-wide baseline, compared to only 5-10% on the 3-prime end.

The excess reduction is fully explained when stratifying by GC content, showing that this is driven by strong GC-biased gene conversion in promoter regions. GC-biased gene conversion acts by preferentially resolving mismatching nucleotides within gene conversion tracts towards G or C bases, therefore reducing diversity similarly as in positive selection but without the broad linkage signature associated with classical selective sweeps. This also causes mutations towards G or C to spread more rapidly within populations (Duret and Galtier 2009). The localisation to promoter regions in canids is likely due to their lack of a functional *PRDM9* gene which, in other mammalian species, localises recombination hotspots away from promoter regions and to *PRDM9*-recognised sequence motifs (Myers et al. 2010). Instead, hotspots are localised in GC-rich promoter regions in canids, driving GC bias and impacting the underlying genealogies (Oliver et al. 2009).

### Boosting power of cross-population selection scans

We next aim to quantify loci of excess differentiation between dogs and grey wolves, which are likely enriched for true candidates under positive selection in early dogs. Recent work has shown that ascertaining for recent coalescences boosts the statistical power of genome-wide F-statistics, an approach implemented in Twigstats (Speidel et al. 2024). We build on this idea to directly compute F_ST_ in genomic windows on inferred genome-wide genealogies and apply a time cutoff that allows us to ignore all coalescences that are older than this cutoff (**Methods**). We thereby develop a new cross-population selection scan, TwigScan, that implements this strategy and improves the statistical power of traditional F_ST_ and f_2_ selection scans that search the genome for excess differentiation between populations, as well as related scans such as the Population Branch Statistic (PBS) (Yi et al. 2010) and its F-statistics equivalent, f_3_ scans.

We propose two mechanisms that improve TwigScan’s ability to highlight selected loci (Supplementary Information). In Twigstats, restricting analyses to recent coalescences reduces standard errors because older coalescences do not directly tag recent admixture events (Speidel et al. 2024). Analogously in TwigScan, by excluding older coalescences, such as those long predating a population split, we remove drift that is uninformative with respect to a recent, population specific selective sweep. A second, distinct mechanism arises from the fact that we only consider a single locus (or limited short genomic windows) and, therefore, few lineages remain as we traverse upwards in a genealogical tree. As a result, the effective sample size at the top of these genealogies is small. The high stochasticity of both the descendants’ make-up of these ancient lineages and their longer branch lengths can contribute substantially to measures of genetic drift between populations, which we are also able to remove in TwigScan (**Figure 3a**).

We used simulations and human data to demonstrate that this approach yields significant power improvements. Across simulations of recent positive selection, where traditional F_ST_ scans are unable to reliably pick out selective sweeps even when selection is strong, TwigScan correctly localised the signal to the sweep region (**Figure 3b**). Applied to 1000 Genomes Project human data (1000 Genomes Project Consortium et al. 2015), well-known loci relating to lactose tolerance and skin pigmentation appeared as outlying regions (*e.g. LCT, SLC24A5, HERC2*) (Fan et al. 2016), while a locus previously reported to be selected for in West Africa (e.g. *SYT1/PAWR*) (Voight et al. 2006) also appeared as outlying (TwigScan F_ST_ z-score of 5.9), whereas traditional F_ST_ scans are unable to identify these peaks of recent excess differentiation (**Figure 3c**, Supplementary Table 4). A comparison to XP-EHH (Sabeti et al. 2007), which detects differences in extended haplotype homozygosity across populations, showed that regions identified by TwigScan also fall in the tail of the XP-EHH distribution, although TwigScan explains only 25–50% of the variance in XP-EHH and so captures distinct, potentially older signatures less strongly characterised by long extended haplotypes or strong selection on *de novo* mutation (SI Figure 8).

### Evidence of strong adaptive introgression in dogs

We applied TwigScan to dogs and grey wolves (**Figure 4a**, **Methods**) by computing TwigScan F_ST_ in 50kbp windows between dogs and Eurasian wolves in both the 722g and Dog10K datasets, applying a time cutoff at 14,000 generations (approx. 40k years ago), predating the grey wolf-dog split. We confirmed in simulations that these parameters are expected to improve power in dog-like demographic histories particularly for selection in the last 9,000 years (Supplementary Information, SI Figures 9, 10). We also conducted a neutral simulation broadly emulating dog and wolf demographic history, which showed that a z-score exceeding six is very rarely achieved genome-wide (Supplementary Information, SI Figure 11).

We therefore highlighted all outlying regions with a z-score greater than six, which identified nine genomic regions (Supplementary Table 5). Of these, several appear as outliers in previous scans based on a mixture of F_ST_, XP-CLR, or Tajima’s D (Pendleton et al. 2018; Freedman et al. 2016; Cagan and Blass 2016). Importantly, all nine regions were also in the tail of TwigScan F_ST_ in the Dog10K dataset (empirical p-value <0.05), which comprises non-overlapping samples relative to the 722g dataset. Among these, a peak on chr6, which includes *AMY2B* and *RNPC3*, and on chr26, which includes *MED13L,* are particularly strongly replicated in the Dog10K genealogies (Supplementary Table 5). These two loci can also be detected in an XP-EHH scan (Zhong et al. 2011) (SI Figure 12), although peaks are less pronounced suggesting increased power in TwigScan in these cases.

The locus with the highest TwigScan score is at the *RNPC3* gene, which lies immediately adjacent to *AMY2B* of which dogs have experienced an increase in copy numbers likely linked to enhanced starch digestion through human contact (Axelsson et al. 2013; Maja Arendt et al. 2014; M. Arendt et al. 2016). While the region has repeatedly been identified in selection scans (Axelsson et al. 2013; Cagan and Blass 2016; Pendleton et al. 2018), the focal genealogy at the *RNPC3* locus further reveals a clear history of introgression with the contributing lineage diverging approximately 3 million years ago from grey wolves (**Figure 4b**). We next conducted several complementary analyses to confirm that this locus is indeed introgressed. Comparing grey wolf-polarised genotypes in multiple extant wolf-like canids revealed that a version of the haplotype is also present in African golden wolves, but is not observed in any other extant wolf-like canid species with available sequencing data (**Figure 4f**). To investigate how diverged this haplotype carried by dogs and African golden wolves is, we computed local f_2_-statistics between species within a genomic window around *RNPC3* (Supplementary Information), which does not rely on genealogies. Strikingly, the *RNPC3* region has the most extreme divergence of either dogs and African golden wolves to other wolf-like canid species across the genome (SI Figure 13a). The pairwise relationships among other canids are not unusual, suggesting no inflation of divergence estimates e.g. due to locally increased mutation rates around *RNPC3*. This suggests also that the introgressing canid is currently unknown and that introgression impacted dogs and African golden wolves only among available extant species. This excess divergence persists even when computed to distant canids such as Ethiopian wolves, dholes, or African wild dogs, which would place the introgressing lineage as potentially more than two million years diverged from dogs (Perri et al. 2021). It is currently unclear if the haplotype introgressed independently into dogs and African golden wolves or if gene flow transmitted the haplotype across these species. Notably, the haplotype in African golden wolves appears slightly divergent to that of dogs (**Figure 4f**).

We next defined the introgressed tract by identifying 305 mutations in perfect linkage disequilibrium (LD) (Supplementary Information, **Figure 4c**). The LD block covers precisely the span of *RNPC3*, and, while our resolution around the immediately adjacent *AMY2B* region is limited due to the low number of SNPs, we do observe some elevated LD upstream of *AMY2B*. However, LD alone is insufficient to map the extent of the introgressed haplotype as the structural complexity of this locus is unresolved in the currently available reference genomes.

To further examine the relationship between introgressed haplotype carrier status and *AMY2B* copy numbers, we assembled a genomic time series spanning the past 16,000 years in Western Eurasia, using 40 ancient and 31 modern dogs (Supplementary Table 6) and quantified haplotype carrier status, as well as *AMY2B* copy numbers, in each individual. While individual SNPs typically only have low sequencing depth in these ancient genomes, we combined transversion SNPs in perfect LD within the *RNPC3* gene to achieve an average 196-fold coverage and make diploid calls for this introgressed haplotype (Supplementary Information). We then inferred copy number variation in modern and ancient canids by aligning to a high-quality Greenland wolf assembly (Sinding et al. 2021) (Supplementary Information).

Interestingly, we notice that four out of five African golden wolves do not carry an increased copy number of the *AMY2B* gene despite at least partially carrying the introgressed haplotype at *RNPC3* (**Figure 4d**, **Methods**, SI Figure 14). Only one African golden wolf from Egypt carries a moderate copy number increase. This individual was previously reported to be admixed with southwest Asian grey wolves and dogs (Gopalakrishnan et al. 2018). In ancient dogs, we observe a significant correlation between haplotype carrier status and *AMY2B* copy numbers (r = 0.68, p = 1.5 × 10⁻⁶). However, several Neolithic Western Eurasian dogs carry the introgressed haplotype in diploid form while lacking any detectable copy number increase at *AMY2B* (SI Figure 14). Conversely, all dogs with clear evidence of an *AMY2B* copy number expansion carry the introgressed haplotype. No such expansions are observed among dogs lacking the introgressed haplotype. This therefore suggests that copy number expansion arose on this introgressed haplotype in ancestors of dogs.

To infer the timing and strength of selection acting on the *RNPC3* introgressed haplotype in dogs, we used CLUES2 (Vaughn and Nielsen 2024) to infer an allele frequency trajectory; this modelling suggests a selection coefficient of 0.003 (p = 2.6 x 10^-8^), with onset of selection dating approximately to 8,000 years ago (**Figure 4e**). This coincides with increased farming practices among humans in Western Eurasia and therefore likely increased exposure to starch rich diets in dogs. This estimate is also consistent with a previous study that detected high copy-number variation of *AMY2B* in dogs from Romania and Serbia 7,000 years ago (Ollivier et al. 2016; Bergström et al. 2020). In our analysis (SI Figure 14), the oldest dog with a suggestive increase is a 6.8k year-old low-coverage dog from Serbia who carries the introgressed haplotype at heterozygous status, and the oldest dog with a copy-number increase matching modern dogs is a 5.8k year-old dog from Iran. However, several Neolithic dogs carry the introgressed haplotype but show no copy number increase, suggesting regional stratification at this time (SI Figure 14). It is therefore unclear whether the haplotype initially conferred a separate advantage, or if it was swept to near fixation only after *AMY2B* duplicated on its background. Today, the derived allele is found across most dog breeds but is absent in some arctic breeds and dingoes (**Figure 4f**).

Applying this framework to the *EPAS1* locus, we can further replicate adaptive introgression in Tibetan dogs and wolves associated with high-altitude adaptation, which has previously been reported as introgressed from a yet uncharacterised canid population (Witt and Huerta-Sánchez 2019; M.-S. Wang et al. 2020; Gou et al. 2014) (SI Figure 13d). Computing local f_2_-statistics for this region suggests that the unknown canid is basal to golden jackals, but less diverged compared to the ghost source responsible for the *RNPC3*/*AMY2B* haplotype (SI Figure 13c).

### Recent adaptation in dogs

We next applied a single-population selection scan that is also able to condition on a specific time period in a genealogical history (**Methods**). We used this test to detect more recent rapid adaptation in dogs focussing on the last 10,000 years, post-dating initial domestication and may reflect more localised instances of dog adaptation.

Our test is an adaptation of the Relate selection test (RST) (Speidel et al. 2019), which computes the probability that a mutation has outcompeted other lineages to spread to at least its present-day observed frequency, conditioning on the number of lineages (k) present when the mutation arose. This test uses the distribution on partitions implied by the standard coalescent, which is unaffected by demographic change in the genealogy (Griffiths and Tavaré 1998). Here, we use a simple extension of this test (Relate selection test over epochs; RST-Over Epochs), which counts the number of lineages that carry the derived and ancestral alleles at a pre-specified time, and computes the probability that the derived-allele-carrying lineages subsequently expanded to at least the present-day observed frequency.

We demonstrate the increased power (smaller p-values) of the modified RST-Over Epochs test for selection under standing variation, where the power gain increases with age of the mutation under selection (SI Figure 15, Supplementary Information). For selection acting throughout the lifetime of a mutation, as expected, we observe no increase in power. However, we note that SNPs linked to *de novo* mutations under selection experience increases in frequency that resemble selection on standing variation, and this test therefore is also expected to highlight the broader haplotypic background of a positively selected mutation.

We applied this test to dogs in the 722g dataset, conditioning on the mutation trajectories over the last 10k years. We identify three peaks from this timepoint (chr19, chr26 and chr8), all of which show broader LD peaks than when computed across the lifetime of the mutation (**Figure 5a**, SI Figure 16). We dismiss the peak on chromosome 8 given its proximity to the end of the chromosome. The peaks on chr19 and chr26 are replicated in an iHS scan (normalised iHS value > 6, SI Figure 12). Interestingly, they appear driven by different dog populations. The peak on chromosome 26 falls within a cluster of immunoglobulin mRNAs and appears to be driven by rapid recent coalescence events in Western dogs (**Figure 5d**, SI Figure 17). In contrast, the signature on chr19 appears to be more widespread amongst dog populations (**Figure 5c**, SI Figure 18). This signature falls amongst a number of uncharacterised transcripts and, notably, is 200kbp downstream from the aligned *H. sapiens GPR148* gene, predicted to play a significant role in olfaction.

In both peaks, candidate SNPs (SNPs within 400Kb of the top-ranking SNP of the chromosome -log₁₀(p) > 7) are recorded with higher evidence of selection within the last 10kya than over the lifetime of the mutation. This is particularly striking for SNPs in the peak of chr19, which have log₁₀(p) difference of up to 10 when compared to the lifetime p-value. Further, both peaks contain SNPs with low selection p-values that are at intermediate or high frequency in wolves (SI Figure 19), suggesting positive selection in the last ∼10k years has driven rapid increases in the frequency of old, segregating variation.

## Discussion

By inferring joint genealogies for hundreds of canids, our analysis identified several unique evolutionary signatures shaping the canine genome. Our genealogies were able to recapitulate dog and grey wolf demographic history consistent with the literature (Skoglund et al. 2015; Bergström et al. 2022), highlight a species-wide sweep at *IFT88* identified previously using ancient DNA (Bergström et al. 2022), and show the expected effects of purifying selection around genes (Hernandez et al. 2011), but with an additional strong effect of GC-biased gene conversion acting on gene promoters. We further identified mutation rate pulses which were verified using independent mutation count ratios, and adaptive introgression at *AMY2B*/*RNPC3* which was verified directly using genotype divergence relative to other wolf-like canids, copy number estimation, and ancient DNA time series modelling. We were able to replicate our results across two independent datasets and demonstrate the potential of genealogies to jointly capture a wide range of evolutionary processes shaping the canine genome.

At present, the cause for the two identified strong, recent mutation rate pulses private to dogs remain unknown, and available de-novo mutation data in dogs is inconclusive as to whether these pulses remain present in dogs today. Ancient DNA shows that the dog-wide mutation rate pulse in the context CA/CT is already observed in ancient dogs 10,000 years ago, predating the formation of recent breeds. A similarly strong mutation rate pulse in a different nucleotide context was previously reported in humans, traceable to 10k - 30k years ago and primarily impacting populations from Western Eurasia and South Asia, with potential origins in ancestors of early farmers (K. Harris 2015; Kelley Harris and Pritchard 2017; Speidel et al. 2021; DeWitt et al. 2021). The pulse in humans is therefore relatively localised in time and space and arose likely in southwest Asia following the out-of-Africa bottleneck, suggesting a genetic cause or a tightly constrained environmental cause. If the pulse in dogs is genetic in origin, reduced effective population sizes in dogs may have weakened selection against mutagenic variants.

Notably, we do not identify comparable pulses in grey wolves. Alternatively, if environmentally caused, these pulses may reflect environmental influences associated with close association with humans. Regardless of the mechanism, our results indicate that germline mutation rates can be highly dynamic on shorter evolutionary timescales within specific mutational contexts.

Our genealogies further capture strong signatures of selection around genes, including widespread reduction in the TMRCA consistent with purifying selection, GC-biased gene conversion, and species-wide selective sweeps, such as at the *IFT88* locus. These processes can affect genealogies in similar ways and produce comparable reductions in TMRCA. Distinguishing between species-wide purifying and positive selection using genealogies remains difficult but, by quantifying if lineages experience a persistent reduction in population size or selective-sweep-like dynamics, future work may help resolve these scenarios.

To identify recent adaptation in dogs, we applied a new genealogy-based cross-population selection scan, TwigScan, together with a time-conditioned implementation of the Relate selection test. These approaches identified multiple loci with evidence of positive selection. Among these, the region containing the *RNPC3* and *AMY2B* genes, previously implicated in dietary starch-associated adaptation in dogs (Axelsson et al. 2013), showed evidence of introgression from a deeply divergent canid lineage, followed by a rapid increase in frequency approximately 7,000 - 8,000 years ago. Interestingly, the introgressed haplotype is only incompletely associated with copy number variation at *AMY2B.* In particular, four out of five African golden wolves have no copy number expansion but carry an introgressed haplotype, while some Neolithic dogs from Europe also show no copy number increase despite clear evidence of introgression. It remains possible that the introgressed haplotype conferred a separate earlier advantage, or that it was swept along with later copy number increases occurring on its background. Future work may investigate the detailed relationship between the introgressed haplotype and mechanistic links to *AMY2B* copy number changes. Humans also experienced copy number increases at Amylase genes (Bolognini et al. 2024; Yilmaz et al. 2024), although the original copy number increase appears ancient and the evidence for selection linked to starch rich agricultural diets remains debated (Yilmaz et al. 2024; Soler i Núñez et al. 2026). In dogs, the copy number increase appears more clearly linked to the Neolithic transition, and genealogies reveal a history of adaptive introgression. However, the precise origins of this haplotype, when or how it introgressed into dogs (and African golden wolves), and how copy numbers at *AMY2B* subsequently changed remains unresolved.

The geographic origin and early history of dogs remains debated (Shannon et al. 2015; G.-D. Wang et al. 2016; Savolainen et al. 2002; Bergström et al. 2020; Gojobori et al. 2024). This work and others (e.g. on the *EPAS1* locus (M.-S. Wang et al. 2020)) highlight the possibility that introgression with uncharacterised canid lineages has contributed to dog adaptation – which may reflect the propensity for interbreeding among canids (Gopalakrishnan et al. 2018; Lin et al. 2025). Together, our results show that genealogical inference can capture the complex evolutionary history of dogs and their relationships to broader canid diversity, enabling increasingly detailed reconstructions of early dog evolution.

## Acknowledgements

We are grateful to Guo-Dong Wang, Shuhua Xu, Shufang Jia, Pontus Skoglund, and Aida Andres for helpful discussions. Analyses were conducted using the Francis Crick Institute compute resources and the HOKUSAI supercomputer at RIKEN (project ID RB240043). L.S. acknowledges JSPS KAKENHI grant 24K23946. A.B. was supported by the Leverhulme Trust (grant no. PLP-2023-281). JR thanks Rango for his canine support. For the purpose of open access, the authors have applied a CC BY public copyright licence to any Author Accepted Manuscript version arising from this submission.

## Author Contributions

L.S. conceived the study, L.S. and M.S. led the development and application of TwigScan, L.S. and J.R. led the Relate analysis, L.S. and D.T led the TMRCA analysis. J.R., M.S., D.T., A.C.V., L.S. performed data analysis, J.R., M.S., L.S. wrote the manuscript, all authors edited the manuscript.

## Code availability

TwigScan is freely available within the Twigstats R package under an MIT licence (https://leospeidel.github.io/twigstats/reference/TwigScan.html).

## Software availability

bwa: https://github.com/lh3/BWA; htsbox: https://github.com/lh3/htsbox; Relate v1.2.1 https://myersgroup.github.io/relate/; relate_lib: https://github.com/leospeidel/relate_lib; tskit: https://tskit.dev/software/tskit.html; pyslim:https://github.com/tskit-dev/pyslim; SLiM:https://messerlab.org/slim/; SELSCAN: https://github.com/szpiech/selscan

## Data availability

Reference genome UU_Cfam_GSD_1.0 was downloaded from https://www.ncbi.nlm.nih.gov/datasets/genome/GCF_011100685.1/ and reference genome canfam3.1 was downloaded from https://www.ncbi.nlm.nih.gov/datasets/genome/GCF_000002285.3/. Recombination rates were downloaded from https://github.com/auton1/dog_recomb. Dog10K phased data was downloaded from https://kiddlabshare.med.umich.edu/dog10K/phased-imputation-panel/. The 722g genotypes were obtained from BioProject PRJNA448733. Gff files containing gene annotation for canfam3.1 and canfam4 were downloaded from https://www.ncbi.nlm.nih.gov/datasets/genome/GCF_011100685.1/ and https://www.ncbi.nlm.nih.gov/datasets/genome/GCF_000002285.3/. The Andean fox genome was downloaded from BioSample SAMN02487034. Inferred genealogies, TMRCA, and mutation rate estimates have been deposited to Zenodo (https://zenodo.org/records/20339492).

## Materials and methods

We provide full details in the Supplementary Information and an outline of our analyses below.

### Canid genealogies

We selected samples from the NHGRI Dog Genome Project dataset (722g) (Plassais et al. 2019) and Dog10K dataset (Meadows et al. 2023) as shown in SI Figure 1, 2 and Supplementary Tables 1, 2. The 722g genealogies include 135 canids (95 geographically diverse dogs, 39 grey wolves, and 1 coyote) and genomes are aligned to CanFam3.1. The Dog10K genealogies include 338 canids (281 village dogs and 57 grey wolves) and genomes are aligned to UU_Cfam_GSD_1.0. Further details on genealogy inference are provided in the Supplementary Information. Genealogies, mutation rates, TMRCAs are available in a Zenodo repository (https://zenodo.org/records/20339492).

### Mutation rate inference

We quantify mutation rate pulses in two distinct ways, which we describe in more detail in the Supplementary Information and we provide scripts to replicate these in Zenodo (https://zenodo.org/records/20339492).

First, we quantify mutation rates in samples built into a Relate genealogy. We inferred mutation rates for 96 triplet mutation categories using the function RelateMutationRate – mode ForCategoryForPopForChromosome provided with the Relate package (SI Figure 3). This function can compute mutation rates for subpopulations, which we use to compute separate mutation rates for dogs and grey wolves. We obtain confidence bands using 100 block bootstrap iterations, where each block consists of 1000 consecutive trees. We postprocess these mutation rates to compute time normalised rates, as detailed in the Supplementary Information. We further used the same function to compute mutation rates to quantify GC-biased gene conversion, by defining categories of mutations towards and away from G/C nucleotides, and stratifying by recombination hotness (Supplementary Information).

Second, we quantified mutation rates without using full genealogical information by simply computing the ratio of mutations falling in a specific context (accounting for strand symmetry) and the number of mutations falling into the mutating single nucleotide category, e.g., CA/CT and TG/AG divided by A/T and T/A (Supplementary Information). This approach allowed us to quantify the pulse in ancient genomes that were not built into Relate genealogies (e.g. due to low coverage), in addition to other modern genomes. Samples used for this analysis are listed in Supplementary Table 1 and 3. To improve sensitivity to recent mutation rate pulses unique to dogs (as identified in our Relate analysis), we ascertained for younger mutations whose upper end of the branch is < 10,000 generations (30,000 years) old in the 722g genealogy. We used a block-jackknife to compute standard errors (Supplementary Information). We further verified that the mutation signatures remained when computed only using mutations that were observed in more than one individual (Supplementary Information).

### Species-wide TMRCAs for dogs and wolves

To compute the TMRCAs of all dogs and wolves along the genome, we converted Relate genealogies into tskit format (Kelleher et al. 2016) using the Convert function in the relate_lib package (Software Availability). We then computed a rolling mean TMRCA with window size 50kb, after restricting to bases where >80% of bases within its 1Mb vicinity were passing in a custom mask (Supplementary Information).

We identified regions with the youngest TMRCAs by first sorting the genome-wide TMRCAs by their age. We then chose the youngest TMRCA in this list and excluded 1Mb to each side of this chosen locus before proceeding to choose the next youngest TMRCA. We applied this procedure to the 722g genealogies to identify the loci containing the youngest 20 TMRCAs genome-wide, of which 19 overlapped with a gene, and show the TMRCAs surrounding these loci in both the 722g and Dog10K datasets in SI Figure 7.

### TwigScan

TwigScan takes Relate genealogies as input and computes f_2_ or F_ST_ in windows across the genome, where we use the definition of f_2_ and F_ST_ provided in Ref (Patterson et al. 2012). From these values, one can also compute a PBS statistic by transforming T(A,B) = -log(1-F_ST_(A,B)) and computing PBS(A;B,C) = (T(A,B) + T(A,C) - T(B,C))/2. Similarly, the corresponding statistic given f_2_-statistics is the f_3_-statistic, computed as f_3_(A;B,C) = (f_2_(A,B) + f_2_(A,C) - f_2_(B,C))/2.

The TwigScan implementation is available as part of the Twigstats package (https://leospeidel.github.io/twigstats/reference/TwigScan.html).

We provide intuition for how TwigScan reduces noise in f_2_ or F_ST_ scans in the Supplementary Information.

### Relate selection test

We apply the Relate selection test which is implemented in the Relate package. This function quantifies to what extent lineages carrying a mutation have outcompeted other lineages that do not carry the mutation. This test can be applied in two separate ways: We can either compute the p-value for the lifetime of a mutation (Relate selection test [RST]), or we can compute a p-value that tests for selection since some time *t* in the past ([RST-Over Epoch]) (Supplementary Information).

### Other selection tests

All haplotype-based statistics were calculated using the SELSCAN programme (Szpiech and Hernandez 2014).

### Simulations

We used the forward-simulator SLiM (Haller and Messer 2019) and associated software pyslim (Gopalan et al. 2025) to model several scenarios of positive selection, including under canid demographic history. Specific details are provided in the Supplementary Information.

### *TwigScan* applied to the 1000 Genomes Project data

We applied the *TwigScan* function to genome-wide genealogies inferred by Relate from the combined 1000 Genomes Project and HGDP datasets (Koenig et al. 2024). We computed F_ST_ between the GBR and YRI populations and averaged them over 50kb windows in two different ways. First, we computed conventional F_ST_ using allele frequencies and applied no time cutoff (infinite generations). Second, we used Relate branch lengths directly and applied a 2000 generation (∼56k years) time cutoff. We show these two sets of F_ST_ runs in Figure 3c, and provide a list of outlying regions in Supplementary Table 4 (details in Supplementary Information).

### *TwigScan* applied to the Dog10K and 722g data

We applied TwigScan to the genome-wide genealogies inferred by Relate from the 722g and Dog10K datasets. We computed TwigScan F_ST_ between the dogs and wolves and, as for the human dataset, averaged these over 50kb windows using an infinite time cutoff and a cutoff of 14,000 generation (∼42k years), chosen to exceed the dog-grey wolf split time.

We identified outlying regions in the 722g genealogies with a z-score exceeding six (based on expectations in a neutral simulation, Supplementary Information). For each of these regions we then computed an empirical p-value in the Dog10K dataset, by first lifting over regions using the UCSC liftover tool and then computing the proportion of windows in Dog10K that had at least the observed TwigScan F_ST_ score. These are listed in Supplementary Table 5.

Among the hits we identified was the locus around the *RNPC3* gene, immediately adjacent to *AMY2B*. This region has repeatedly been detected in selection scans (Axelsson et al. 2013; Cagan and Blass 2016; Pendleton et al. 2018). We plotted the local genealogy at *RNPC3* using the TreeView function provided in the Relate package, which revealed a deep 3M year old coalescence time between dogs and grey wolves. We mapped this introgressed haplotype and investigated whether other wolf-like canids carried the haplotype, focussing on coyotes, red wolves, African golden wolves, golden jackals, dholes, Ethiopian wolves, and African wild dogs, using genomes from (Liu et al. 2018; Gopalakrishnan et al. 2018; Plassais et al. 2019; Kardos et al. 2018; Sinding et al. 2018). A locally increased mutation rate could drive this observed deep coalescence between dogs and wolves and to exclude this possibility, we evaluated whether relationships among other canids is unusual. We investigated how the introgressed haplotype relates to copy number variation at *AMY2B,* identifying that *AMY2B* duplications are only observed on the introgressed background. We further modelled the allele frequency trajectory of this haplotype using CLUES2 (Vaughn and Nielsen 2024). We describe these analyses in more detail in the Supplementary Information.

### *Relate* selection test applied to the 722g data

We subset the 722g genealogies to dogs. We then used the Relate function RelateSelection –mode Quality to compute the number of mutations mapping to trees and the proportion of branches with at least one mutation. We then removed approximately 5% of trees with the lowest quality, measured using these two quantities, by applying the filter.allele_ages function in the relater R package. As an additional filtering step, we also removed SNPs with derived allele frequency of less than 5%. This resulted in 7,765,429 SNPs (41.6%) for analysis in dogs.

We then applied the Relate selection test (evaluating the evidence for positive selection over the lifetime of the mutation) and the Selection Over Epochs test (specifically testing for selection in the last 10kya). Manhattan plots comparing both runs are shown in SI Figure 16. We identified SNPs with evidence of positive selection at the genome-wide significance level (p-value < 5e-8). This identified three peaks on chromosome 19 (SI Figure 16), chromosome 26 (SI Figure 17), and a hit on chromosome 8 which fell close to the end of the chromosome and was therefore discarded. These hits were also confirmed using iHS (SI Figure 12).

