## Supplementary Information for "Joint dog and wolf genealogies reveal the evolution of the canine genome"


Sample selection in the 722g and Dog10K datasets 2
Andean fox genome 3
*Relate genealogies* *3*
Additional modern and ancient canid genomes 4
Mutation rate inference 6
  Mutation rates using Relate genealogies 6
  Mutation rates in trinucleotide contexts using Relate 6
  GC-biased gene conversion using Relate 7
  Quantifying mutation pulses in lower coverage genomes 7
  Testing for strand effects and distance from exon 8
  Mutation count enrichment of non-singletons 8
  De-novo mutations 9
Species-wide TMRCAs for dogs and grey wolves 10
The TwigScan method 11
Relate selection test 13
  TwigScan simulations 14
  RST-Over Epochs Simulation 16
TwigScan applied to real data 17
  TwigScan applied to the 1000 Genomes Project data 17
  TwigScan applied to the Dog10K and 722g data 17
Detailed analysis of the AMY2B/RNPC3 locus 18
  Linkage disequilibrium around the AMY2B/RNPC3 locus 18
  Cross-species relationships at RNPC3 and EPAS1 19
  Copy number variation at AMY2B 19
  Allele frequency trajectory at the RNPC3 locus 20
**SI References** **20**
**Supplementary Figures** **26**
  SI Figure 1 Datasets 26
  SI Figure 2 PCA of modern dog samples mapped to canFam3.1 27
  SI Figure 3 Context-dependent mutation rates 28
  SI Figure 4 Context-dependent mutation enrichment in each individual dog sample 29
  SI Figure 5 GC-biased gene conversion 30
  SI Figure 6 Comparison to de-novo mutations 31
  SI Figure 7 Species-wide TMRCAs 32
  SI Figure 8 Comparison of TwigScan and XP-EHH 33
  SI Figure 9: TwigScan performance in simulated dog loci under selection (10kb

windows). 34
SI Figure 10: TwigScan performance in simulated dog loci under selection (50kb windows). 35
SI Figure 11: Comparison of TwigScan FST and standard FST on neutrally simulated canid genomes 36
SI Figure 12 iHS and XP-EHH scans 37
SI Figure 13 Cross species relationships at RNPC3 and EPAS1 38
SI Figure 14 AMY2B copy number variation 39
SI Figure 15 Simulation evaluating the Relate selection test over epochs. 40
SI Figure 16 Relate selection test applied to dogs 41
SI Figure 17 Selection statistics and genealogy for the chr26 peak. 42
SI Figure 18 Selection statistics and genealogy for the chr19 peak. 43
SI Figure 19 Selection evidence in dogs by frequency in wolves 44

### Sample selection in the 722g and Dog10K datasets

We selected a subset of the NHGRI Dog Genome Project (722g) (Plassais et al. 2019) and the Dog10K consortium data (Meadows et al. 2023; Ostrander et al. 2019), and added additional modern and ancient genomes for individual analyses. Supplementary Tables 1 and 2 list samples used for genealogy inference, and Supplementary Table 3 lists additional samples used in this study aligned to the same reference as the 722g dataset (canFam3.1), with columns indicating in which analysis they were used. SI Figure 1 shows the geographic distribution of the modern and ancient dog and grey wolf genomes selected in this study.

For the 722g dataset, we selected individuals from diverse breeds and village dogs following Ref. (Bergström et al. 2022) alongside grey wolves and a coyote, totalling 136 genomes (Supplementary Table 1), ensuring that all samples had a sequencing coverage greater than 8x. Coverages ranged from 8.16x - 66.47x with a median coverage of 18.55x. For the Dog10K dataset, we selected all village dogs and grey wolves, which amounted to 338 genomes (Supplementary Table 2). Coverage ranged from 14x - 29.25x, with a median of 19.7.

The PCA illustrating diversity of dogs in the 722g dataset (SI Figure 2) was performed for all dog samples aligned to canFam3.1 (722g and additional downloaded dog genomes) using the LD-aware gt_pca_autoSVD() and augment() functions in *tidypopgen* (Carter et al. 2026). To maximise the number of individuals represented in the PCA, samples with fewer than 45M (~65%) genotyped positions were excluded from the analysis, and SNPs were filtered out if they were monomorphic or missing in over 10% of remaining samples. Missing genotypes were then imputed using the “mode” method of the gt_impute_simple() function in *tidypopgen*. This resulted in 611,160 SNPs used in the PCA.

#### Andean fox genome

Relate requires an ancestral genome to polarise alleles into ancestral and derived status. We chose the Andean fox as our outgroup. The estimated divergence time between grey wolves and Andean foxes is approximately 4 million years (Chavez et al. 2022), leading to an expected genomic divergence of approximately 1%. No interbreeding between the Andean fox and wolf-like canids is expected, making it a suitable outgroup. We note that Relate tolerates a small amount of misspecified ancestral alleles, e.g. for a mutation observed in dogs/wolves there is approximately a 1% chance that a mutation occurred on the lineage to Andean foxes and the ancestral allele would therefore be incorrect. We obtained an Andean fox genome from Ref (Auton et al. 2013) (PRJNA232497, SAMN02487034). The 722g and Dog10K datasets are aligned to different reference genomes. We therefore first aligned this genome to CanFam3.1 (https://www.ncbi.nlm.nih.gov/datasets/genome/GCF_000002285.3/) and then realigned the genome to reference genome UU_Cfam_GSD_1.0 (https://www.ncbi.nlm.nih.gov/datasets/genome/GCF_011100685.1/). We used bwa mem with default parameters. We then generated a pseudo-haploid genome using htsbox pileup -R -q 20 -Q 30 -l 35 -s 1.

#### *Relate* genealogies

We inferred two sets of genealogies. First, for the 722g dataset, we selected 95 diverse modern dogs, 40 wolves, and 1 coyote. We downloaded an unphased VCF aligned to CanFam3.1 (Data availability). We then used the procedure of the Relate run in (Bergström et al. 2022) to phase the dataset and generate a genome mask file. We used SHAPEIT4 to phase SNPs (Delaneau et al. 2019). Prior to phasing, we first subsetted to samples used in our study (Supplementary Table 1) and then retained only SNPs with less than 1% missingness in genotypes per SNP. We did this as SHAPEIT4 will impute missing genotypes, but these are expected to be of low quality if an excessive proportion of genotypes are missing.

We constructed a genome-wide mask to identify unreliable regions irrespective of whether a variant was observed at a given position. We first applied the GEM mappability tool for k35 (Marco-Sola and Ribeca 2015), excluding regions where short sequencing reads cannot be mapped uniquely. This set ~11.5% of the genome to non-passing. We intersected this mask with the callable regions of the Andean fox genome used as the ancestral genome, which set an additional 0.8% to missing. Finally, we accounted for the fact that the 1% missingness filter applied to the VCF only removed unreliable SNPs. Removing these SNPs without masking the intervening bases could create genomic regions with artificially reduced observed divergence, e.g. if poorly called variants cluster. To identify such regions, we compared one coyote (*Coyote01*) and one Andean fox (*AndeanFox*), two distantly related canids providing enough sequence differences across the genome. We estimated the local density of sites differing between these two individuals before and after excluding SNPs with >1% missingness. Before applying the missingness filter, 99% of 10 kb windows contained more than 50 fox–coyote differences. Regions were masked if, within a rolling 10 kb window, fewer than half of the fox–coyote differences remained after filtering. This filter removed an additional 3.7% of the genome. We downloaded previously inferred recombination rates (Auton et al. 2013). We used a mutation rate of 4e-9 mutations per base per generation and fitted variable effective population sizes using the EstimatePopulationSize function in Relate. This procedure updates the branch lengths in the genealogies to reflect the inferred variable population size history.

Second, for the Dog10K dataset, we downloaded an already phased VCF (Data availability) available through the Dog10K consortium (Data availability). This dataset is available aligned to reference genome UU_Cfam_GSD_1.0 (Data Availability). We selected all 57 grey wolves and 281 village dogs, resulting in a total of 338 samples. We used a per base per generation mutation rate of 4e-9 and a constant recombination rate (1 cM/Mb). We used a genomic mask provided with the data set, which we additionally intersected with the Andean fox ancestral genome. We did not repeat the 1% SNP genotype missingness filter as the data was not phased by us and so this information was not directly available. We used the remapped Andean fox genome as an outgroup to polarise alleles by ancestral state. We then used Relate inferred genealogies for chromosome 1, and jointly fitted time-varying coalescence rates averaged across all samples. We used these average coalescence rate estimates to infer genealogies for all chromosomes 1 to 38.

Ancestral genome, mask files, recombination maps used in our analysis and inferred genealogies are available to download from Zenodo (https://zenodo.org/records/20339492).

To infer human genealogies, we downloaded phased haplotypes from a harmonised version of the 1000 Genomes Project and HGDP datasets (Koenig et al. 2024), selecting individuals of the Yoruba (YRI), Han Chinese (CHB), and British from England and Scotland (GBR) populations. We used Relate input files available from Zenodo (https://zenodo.org/records/15801307) and used a mutation rate of 1.25e-8. We jointly fitted effective population sizes and branch lengths using the EstimatePopulationSize function in Relate.

#### Additional modern and ancient canid genomes

We compiled a dataset comprising all canids listed in Supplementary Table 3, including all samples in the 722g dataset which were not used in the Relate genealogies, several other wolf-like canids obtained from (Kardos et al. 2018; Gopalakrishnan et al. 2018; Liu et al. 2018; Auton et al. 2013; Fan et al. 2016; vonHoldt et al. 2016; Sinding et al. 2018; Wang et al. 2016; Freedman et al. 2014; Perri et al. 2021), as well as 133 ancient canids

(Bergström et al. 2022; 2020; 2026; Marsh et al. 2026; Girdland-Flink et al. 2025; Feuerborn et al. 2021; Sinding et al. 2020; Niemann et al. 2021; Ramos-Madrigal et al. 2021; Mak et al. 2017; Ní Leathlobhair et al. 2018; Lin et al. 2023; Botigué et al. 2017; Frantz et al. 2016; Skoglund et al. 2015; Segawa et al. 2022) (Supplementary Table 3).

Starting from the unphased VCF downloaded for the 722g dataset (data availability), we added additional modern genomes not included in the 722g dataset, using a pipeline described in Bergstrom et al. 2022. In brief, downloaded FASTQ files were mapped to the canFam3.1 reference genome using BWA *mem* with default parameters (Li 2013). Duplicate reads were identified using Picard Tools (https://broadinstitute.github.io/picard/) ("Picard," n.d.). Genotyping was carried out for SNPs identified in the 722g dataset VCF using GATK HaplotypeCaller before being merged into the larger dataset using BCFtools (Danecek et al. 2021). The VCF was then filtered to minimise missingness and very low frequency variants as in Bergstrom et al. 2022, resulting in a set of 67.8M SNPs.

Ancient genomes were processed as described in Bergstrom et al. 2022 for most samples. In brief, paired-end data was collapsed and adapters were trimmed using SeqPrep, filtering out unmerged reads (https://github.com/jstjohn/SeqPrep) (St. John, n.d.). Reads were mapped using BWA aln using aDNA-adapted parameters (-l 16500 -n 0.01 -o 2) (Li and Durbin 2009). Duplicates were identified using orientation and start and end coordinates were removed. For the remaining samples, the nf-core/eager v2.3.3 pipeline was used to process samples in a similar way (Fellows Yates et al. 2021). Differences include using *fastP* v0.20.1 to trim poly-G tails from two-colour chemistry Illumina sequencing data (NovaSeq, NextSeq), using AdapterRemoval v2.3.1 to collapse paired-end data and trim adapters, applying slightly different parameters for BWA aln v0.7.17-r1188 (-o 1 instead of -o 2), removing duplicate reads using DeDup v0.12.8 for paired-end data or Picard MarkDuplicates v2.22.9 for single-end reads, and using DamageProfiler v0.4.9 to calculate terminal 5' deamination rates (Chen et al. 2018; Schubert et al. 2016; Li and Durbin 2009; Peltzer et al. 2016; "Picard," n.d.; Neukamm et al. 2021). Pseudohaploid genotypes were generated for all ancient genomes using htsbox r345 filtering for reads of 35bp or longer and of mapping quality above 20, as well as base qualities above 30 (https://github.com/lh3/htsbox).

We used these additional modern and ancient genomes to reconstruct the temporal trends, and representation across wolf-like canids of the mutation rate pulses we identify, as well as to quantify selection and introgression at the *AMY2B*/*RNPC3* locus associated with starch digestion.

#### Mutation rate inference

We quantify mutation rate pulses in two different ways: we use Relate genealogies to infer mutation rates through time and we use a simpler count of mutations occurring in each nucleotide context. The second approach works in lower coverage ancient genomes which are not suitable to incorporate into Relate genealogies and relies less on the inferred genealogies (other than by pre-ascertaining mutations by age to enrich for more recent mutations). We describe both approaches below. We provide scripts and result files in a Zenodo repository (https://zenodo.org/records/20339493).

**Mutation rates using Relate genealogies**

Given a Relate genealogy, Relate provides a function to infer mutation rates through time. This function infers mutation rates using a simple maximum likelihood approach conditional on genealogies by counting the number of mutations of a certain type in each time period, and dividing by the total opportunity for that mutation to occur (Speidel et al. 2019). The opportunity is computed for each mutation category by multiplying the total branch length in that time period with the number of bases that could have mutated in a given mutation category. We account for strand symmetry. For instance, for a C to T mutation, we count the number of C and G nucleotides in the genome passing the genomic mask, as these had a potential to mutate into a T or A, respectively.

This is implemented in the function RelateMutationRate –mode ForCategoryForPopForChromosome provided with the Relate package. We used arguments --years-per-gen 3, --bins 3,7,0.2, and specified the fox genome as the ancestral genome and used the same genomic mask file used for inference of the Relate genealogies. To obtain uncertainty estimates of rates, we computed 100 block bootstrap iterations, where each block consisted of 1000 consecutive trees. It is further possible to compute mutation rates only using lineages ancestral to a subset of samples in the genealogy. We used this option, implemented as the argument --pop-of-interest, to compute separate mutation rates for dogs and grey wolves.

**Mutation rates in trinucleotide contexts using Relate**

We investigated how mutation rates depend on their upstream and downstream nucleotide context, by computing triplet mutation rates, e.g. ACG mutating towards AGG. Strand symmetry implies that e.g., the motif CGT mutating towards CCT on the forward strand would correspond to ACG mutating towards AGG on the reverse strand. This is computed using the argument –mutcat in the function above, grouping triplet mutations by strand symmetry. After accounting for such strand symmetries there are 96 different triplet mutation categories. We inferred the mutation rates through time for each of these triplet categories (SI Figure 3).

To identify mutation pulses, we postprocessed the inferred rates as follows. The average mutation rate across all categories is confounded by variations in the effective population size and so any fluctuations in the average mutation rate cannot be distinguished from imperfectly inferred effective population sizes. Therefore, for each population, we normalised the mutation rate across mutation categories by first calculating in each time period the mean mutation rate across all different categories. We then divided the inferred mutation rates by this mean mutation rate. For each mutation category, we further normalised the mutation rate across time, by computing the mean across log-spaced time epochs, and dividing by this mean.

**GC-biased gene conversion using Relate**

In the 722g dataset, where we had access to a previously inferred recombination map, we inferred mutation rates for each of the six single nucleotide substitutions, stratified by recombination hotness (SI Figure 5). We used the recombination map obtained from (Auton et al. 2013). We did not repeat this for the Dog10K dataset as no recombination map is currently available for this reference genome.

We defined four levels of hotness: we defined recombination rates in the bottom 5th percentile as ‘very cold’, recombination rates between 5th and 10th percentiles as ‘cold’, between 10th and 95th percentiles as ‘medium’, and greater than the 95th percentile as ‘hot’. We computed mutation rates for these categories by amending the genomic mask file that we supply to the function RelateMutationRate –mode ForCategoryForPopForChromosome, setting regions artificially as non-passing if they do not fall within the recombination hotness of interest.

For each hotness level, we normalised the mutation rates across all six nucleotide substitutions and across time as before and examined differences in mutations mutating towards or away from G/C.

**Quantifying mutation pulses in lower coverage genomes**

We quantified mutation rates in ancient and modern genomes, many of which were of low coverage and therefore not suitable in the Relate analysis. These are listed in Supplementary Table 3. We used the VCF compiled as described in section “Additional modern and ancient canid genomes”. As the pulse is expected to be recent, to enrich for mutations associated with the pulse, we ascertained SNPs within this VCF that were dated in the 722g genealogies (see Zenodo for inferred genealogy). We excluded SNPs that did not map to the genealogy (is_not_mapping = 1) or were flagged as flipped (is_flipped = 1), i.e. the ancestral and derived alleles were flipped in order to map the mutation to the genealogy. We then chose to retain SNPs where the upper end of the branch is <10,000 generations (30,000 years) old. We then computed the proportion of mutations falling in a specific context by dividing the counts for that context (accounting for strand symmetry) by the number of mutations falling into the mutating single nucleotide category, e.g., CA/CT and TG/AG divided by A/T and T/A. Standard errors were computed by defining blocks of size 10Mb, and computing a block jackknife.

**Testing for strand effects and distance from exon**

For mutations <30k years old (estimated by the midpoint of a branch), we binned these by distance to the nearest exon (SI Figure 3). For each bin, we computed the proportion of mutations in a given trinucleotide context (not accounting for strand symmetry here as we wanted to test if the same pattern holds for each strand-corresponding motif), relative to the total number of mutations. We did this separately for mutations that are unique to dogs (or African dogs) $p_{unique}$ and other mutations $p_{not\_unique}$. We then computed the fold enrichment as the ratio $p_{unique}/p_{not\_unique}$. This enrichment is therefore quantifying whether the proportion of mutations of that type is increased in dogs, relative to mutations not unique to dogs. We used a weighted block jackknife over chromosomes to obtain confidence intervals.

As a control, we used CpG transitions. While these have an elevated mutation rate relative to other mutation types, we do not necessarily expect an effect by distance from the nearest exon, and we do not expect CpG transitions to be particularly enriched in dogs. As expected, the fold-enrichment of CpG transitions unique to dogs is close to 1, regardless of distance to exons (SI Figure 3e).

For GC/G and CA/T mutations, we overall see an enrichment of mutations unique to dogs, as expected. However, distance from exon appears to have little effect overall, with the exception of GCG mutating towards GGG. These appear to be depleted closer to exons (SI Figure 3c,d).

**Mutation count enrichment of non-singletons**

To further rule out potential sequencing artefacts, we quantified whether mutation enrichment is observed after restricting to non-singletons (SI Figure 4). Specifically, in addition to using only SNPs with an upper age <10,000 generations as before, we restricted our analysis to SNPs for which the derived allele was carried by at least two individuals. To do this, we selected individuals listed in SI Table 1 and SI Table 3 with column “canid_type” set to one of "grey_wolf", "breed_dog", "dog", "free-ranging/indigenous_dog" and age equal to 0, excluding one individual with analysis_id “1185”, as this sample was added later. We then required at least two individuals to carry the mutation in either heterozygous or homozygous derived-allele status, polarised relative to the Andean fox. Any artefact impacting one individual, and therefore causing a spurious SNP, would therefore be excluded from our analysis.

For each dog sample, we counted the number of mutations within each of the 96 triplet mutation types relative to the corresponding count in grey wolves. For each sample, we then normalised these enrichments to 1 by dividing by the median enrichment across all mutation categories. We used a block jackknife with block size 50Mb to quantify uncertainty.

The resulting enrichment is in agreement with our Relate-inferred mutation rate estimates (SI Figure 4). Given that the mutation rate pulses estimated directly from Relate genealogies replicate across both the 722g and Dog10K datasets, which contain distinct samples aligned to distinct reference genomes, and given that the signatures also replicate when quantified as a mutation count enrichment, including when we use exclusively mutations carried by multiple individuals, and also show strand symmetry as expected, we therefore conclude that these signatures are unlikely to be artefacts.

**De-novo mutations**

We used de-novo mutation counts provided by Ref. (Zhang et al. 2025), who sequenced 390 trios from 43 breeds (SI Figure 6). Sample sizes were too small to consider triplet mutations and we therefore focussed on stratifying mutation categories just by one upstream or one downstream nucleotide. We did this for non-African dog breeds and for Basenji dogs separately, expecting different patterns given the identified mutation rate pulses in these groups.

For a visual comparison, we used Relate-inferred mutation rates (--bins parameter 2,6,0.2), where we initially computed mutation rates for all 96 triplet categories. As before, we removed average temporal trends shared across all mutation categories by dividing by the category-specific mean rate in each epoch. We then averaged these to only stratify by one upstream (or downstream) nucleotide. We additionally averaged over epochs (between $10^3$ and $10^4$ years ago) to obtain a single mutation rate estimate for each category and normalised these relative rates to sum to 1, to compute the expected proportion of mutations in each mutation category.

This indicated, for instance, that in grey wolves, we expect approximately similar numbers of mutations of types CA/CT, CC/CG, and CA/CC, while in dogs, the category CA/CT is expected to be elevated. In observed de-novo mutation counts, we did not observe an increased count for CA/CT relative to other categories, indicating that the mutation pulse is no longer active in these modern-day dogs.

We also computed explicit multinomial likelihoods, which can be used to test if the observed de-novo mutations are more likely to originate from a grey wolf-like or dog-like mutation spectrum. However, these appeared unreliable, for instance, while across all categories it appears that observed de-novo mutation counts are more likely to originate from dogs, this result reversed once we excluded C/T mutations which dominate the count. Overall, the ranking across different mutation categories does not appear to support the existence of the mutation rate pulse in these dogs.

For the Basenji dog, we relied on data of six trios, likely not enough to make confident conclusions. However, interestingly, we observed no de-novo mutations in categories CA/CT and GC/GG, which we expect to be enriched if mutation pulses were active.

#### Species-wide TMRCAs for dogs and grey wolves

We describe how we computed species-wide TMRCAs for dogs and grey wolves. We first converted Relate trees to tskit format (Kelleher et al. 2016) using the Convert function in the relate_lib package (https://github.com/leospeidel/relate_lib). For the 722g genealogies, we then computed the TMRCA after first simplifying the tree sequence to exclude the coyote, while for Dog10K, which did not include a coyote, we simply computed the root TMRCA of the genealogies.

To ensure that TMRCAs were reliably inferred by Relate, we computed a new genomic mask file where we required sufficient data in the 1Mb vicinity of a base pair. We filtered TMRCAs using a rolling mean genomic mask, compiled for each dataset as follows. We started with the genomic masks used in Relate inference described above, each provided by the Dog10K project, and the 722g dataset. We recoded each mask such that a ‘passing’ base-pair is recorded as 1 and a ‘non-passing’ base-pair is recorded as 0 and computed a rolling mean with window size of 1Mb. We set any base-pair with a value of less than 0.8 as ‘non-passing’ in addition to the already non-passing base-pairs, so any passing base-pair now has at least 80% of bases passing in its 1Mb vicinity. We then computed a rolling mean of this mask, again with a window size of 1Mb, set any values <0.8 as additional ‘non-passing’ bases and repeated this procedure for a total of 5 times.

We next set any TMRCA at a base-pair that is ‘non-passing’ in the rolling mean genomic mask to be missing. We computed a rolling mean of the remaining TMRCAs using a window size of 50kb. For efficiency, we stored TMRCAs at every 100 bases. In this final dataset, we had TMRCAs for 89.8% and 76.6% of the genome in the Dog10K and 722g project datasets, respectively. We identified the top 20 youngest TMRCAs in the genome by first sorting this list of genome-wide TMRCAs, choosing the youngest, and excluding 1Mb to either side before selecting the next youngest region.

We downloaded genome annotation files (gff files) for each reference genome (data availability) to obtain gene annotations, which we subset to protein-coding genes. We then annotated the top 20 youngest TMRCAs in the 722g genealogies (Figure 2), and identified the corresponding genes in the Dog10K genealogies for comparison (SI Figure 7). Within the 20 regions with the youngest TMRCA, 19 regions have a clear overlap with genes. Around 40-45% of the genome is expected to fall within genes and so 19/20 regions appears to be a significant enrichment. To test this more formally, we used the base-pair with the smallest TMRCA in each of the 20 regions, of which indeed 19/20 cases fell within a protein coding gene, and compared this to 1000 randomly chosen base-pairs passing the TMRCA mask. Of these 441 base-pairs overlapped with a gene, and a two-sided Fisher’s exact test yielded a p-value of 2.5e-06.

We next found that of the 20 regions identified for the 722g genealogies, seven (*ZNF407*, *UVRAG*, *RIMS2*, *NCOA2*, *IL6ST*, *KAT6A*, *RALGAPA1*) had a probability of loss-of-function intolerance score (pLI) of 1 in humans (gnomadv4.1) (Chen et al. 2024). We compared the proportion of these genes (7/19) to the proportion of genes with pLI > 0.99 in gnomadv4.1 (2,832 /18,746) and showed that this is a significant enrichment (Two-sided Fisher's exact test p-value = 0.01703).

Finally, we also computed average TMRCAs by distance to genes, accounting for strandedness (Figure 2, SI Figure 7). We stratified TMRCAs by local GC content. We first computed, for each base in the genome, the proportion of bases in its 1000 base-pair vicinity that are G or C. Given this local GC density estimate, we then grouped bases by quantile ([0,0.5), [0.5,1]), and computed as a function of distance from genes, the average TMRCA in each quantile.

### The TwigScan method

For genome-wide f-statistics, we previously implemented Twigstats, which computes $f_2$-statistics directly on inferred genealogies and ascertains for younger coalescences that tag recent admixture events. Here, we extend this idea to compute $f_2$ or $F_{ST}$ in windows along the genome, where window lengths may be specified in base-pairs, centi-Morgans, or we can compute these for each tree in turn. We use the definitions of $f_2$ and $F_{ST}$ given in Appendix A of Ref (Patterson et al. 2012). We provide documentation for our TwigScan function online (https://leospeidel.github.io/twigstats/reference/TwigScan.html).

We briefly provide two arguments to illustrate why we believe the TwigScan approach will improve our statistical ability to detect positive selection that produces excess differentiation between two populations.

We suppose we compute an $f_2$-statistic between two populations A and B. Let us denote by $f_2(A,B;t)$ the $f_2$-statistic with a time cutoff of $t$. We compute this f-statistic directly on genealogies, and we define $l_b$ to be the expected number of mutations on branch $b$, which is proportional to its branch length and the region in the genome for which the branch persists. Let us suppose that $W$ is the length of the genomic region we consider. Then the $f_2$-statistic can be computed as

$$f_2(A,B;t) \propto \sum_{b\ in\ W} l_{b,t}\,(p_A(b) - p_B(b))^2$$

where the sum goes over all branches $b$ that exist in genomic window W (potentially across several different trees), $t$ is the time cutoff, $l_{b,t}$ is the expected number of mutations on branch $b$ younger than $t$, and $p_X(b)$ is the proportion of descendants of branch $b$ in population $X$. A large $f_2$-statistic is achieved by summing branches with large $l_{b,t}$ and those that are very population specific, such that e.g., $p_A(b) > p_B(b)$.

The first intuitive argument for setting a time cutoff $t$ is that we are interested in quantifying recent selection, and our strategy here excludes drift older than time $t$. Therefore, $f_2(A, B; t)$ will be only influenced by young branches that clearly separate populations A and B, which are good candidates for recent selection in one of the two populations.

The second argument is that older coalescences can contribute substantial variance to an $f_2$-statistic. This is because at the top of the tree, only few lineages remain, making effective sample sizes small. Under the neutral coalescent, for an older branch $b$, $l_b$ can therefore be highly variable; if only two lineages remain, the expected time to coalescence is on the order of the effective population size $2N_e$ with variance $4N_e^2$. At the same time, older branches also tend to have more descendants and so even a random difference in the proportion of descendants from populations A and B can produce a contribution $l_b\,(p_A(b) - p_B(b))^2$ that is comparable to a contribution produced by a branch tagging a recent sweep.

To provide a concrete example, let us consider a short genomic window $W$ that only spans a single tree. Then $l_b$ is proportional to the branch length (as the genomic span of all branches is given by W). Let us assume that the dog-grey wolf split time is 10,000 generations and the effective population size is 50,000 across all populations. Let us first consider a full selective sweep in dogs. If we assume that all dogs coalesce instantaneously, then it will take 10,000 generations for the branch to enter the wolf population. This dog lineage can then coalesce with any wolf lineage remaining at that time and so may coalesce relatively soon with a wolf lineage. The branch tagging the sweep may therefore contribute on the order of $l_b(p_{dog}(b) - p_{wolf}(b))^2 \approx 10000\,(1 - 0)^2 = 10000$. Next, in a neutral region, the contribution of a branch at the top of the tree is $l_b(p_{dog}(b) - p_{wolf}(b))^2$, where $l_b$ has expected length on the order of $2N_e$ with variance $4N_e^2$. In this case, $E[l_b] \approx 100{,}000$. Therefore, as soon as $(p_{dog}(b) - p_{wolf}(b))^2 \approx 0.1$, this neutral branch contributes to $f_2(A, B; t = \infty)$ on a similar order of magnitude to a branch tagging recent strong selection. Because the upper part of the tree is highly stochastic under the neutral coalescent, these large contributions vary substantially across loci and therefore generate background variance in the local $f_2$-statistics.

If the genomic window W spans multiple local trees, then individual older ancestral branches tend to persist over shorter genomic intervals because they have had more opportunity to be broken up by recombination. As a result, the contribution of any single old branch starts to become diluted across the window, and the variance from old coalescences may average out across trees. However, the general principle would still hold: a recent selected branch may span the entire window W, generating a coherent large contribution to $f_2$ across the window, while the collection of older “top-of-the-tree” branches have the potential to produce contributions $l_b(p_{dog}(b) - p_{wolf}(b))^2$ that are comparable in scale.

Overall, excluding the top of the tree therefore focusses the statistic on a relevant recent time period and improves the signal to noise ratio by removing contributions generated by random neutral coalescences at the top of the tree.

### Relate selection test

The Relate selection test computes a p-value that quantifies whether a lineage has more descendants than expected under the standard coalescent model. We use a classical result in coalescent theory that provides a distribution on the shape of coalescent trees (Griffiths and Tavaré 1998). The shape of coalescent trees is invariant with respect to population size changes, and therefore our test is in principle robust to misspecified effective population sizes.

The test can be applied to test for selection either over the lifetime of a mutation or conditional on some time *t* in the past.

We compute the p-value as follows. We first count the number of lineages remaining in the tree at time $t$, which we denote by $k_{all}$. If we are testing over the lifetime of a mutation, we use $t = t_{lower}$, where $t_{lower}$ is the lower end of the branch onto which the mutation is mapping.

Next, we also count the number of lineages at time $t$ carrying the mutation, which we denote by $k_{derived}$. When testing over the lifetime of the mutation and $t = t_{lower}$, then $k_{derived} = 2$.

Finally, we count the number of observed samples (haplotypes) in the present-day carrying the derived allele, which we denote by $n_{derived}$. We denote the total number of sampled haplotypes by $n_{all}$.

The test works by computing how likely it is that $k_{derived}$ out of $k_{all}$ lineages spread to $n_{derived}$ out of $n_{all}$ lineages. In the standard coalescent model, without selection, it is known that $n_{all}$ lineages will be uniformly partitioned into $k_{all}$ ancestral lineages (Griffiths and Tavaré 1998). This result holds irrespective of demographic history, and this is then used to compute a p-value that at least $n_{derived}$ haplotypes subtend from $k_{derived}$ ancestral lineages (Speidel et al. 2019).

### Simulations

#### TwigScan simulations

To evaluate TwigScan, we simulated two populations that split 3500 generations ago and inserted a positively selected mutation into one of these two populations. To achieve this, we used SLiM to simulate two populations with constant diploid population size of 10,000 for 3500 generations. While the aim was to keep parameters simple, they were chosen here to broadly emulate human-like demography, with a split time on the order of the earliest estimates of out-of-Africa dispersals. We simulated 10Mb genomes with a recombination rate of 5e-9 events per base per generation and no mutations. In generation 2,750 of this simulation, we inserted a mutation at the centre of a randomly selected genome in population 1 and assigned a haploid selection coefficient of 1% to this mutation. If the mutation was lost, we restarted the simulation at time point 2,750. Once the mutation reached 80% frequency, we changed the selection coefficient to 0.

We then used pyslim (Gopalan et al. 2025) to recapitate this simulation. We chose an ancestral population size of 10,000 and a recombination rate of 5e-9. We then randomly sampled 100 diploid individuals from each population and added mutations with a rate of 1.25e-8 events per base per generation. We output the true tree sequence, as well as a vcf of the 400 sampled genomes.

Given these data, we inferred Relate trees with a constant haploid population size of 20,000 and mutation rate of 1.25e-8. We also converted the true tree sequences to Relate format using the relate_lib package. Finally, we computed $F_{ST}$ between the two populations using the function `Fst_blocks_from_Relate`, with a bin size of 10,000 bases and option 'dump_blockpos' enabled which records the base pair position of each block. As shown in Figure 3b, applying a TwigScan cutoff of 2000 generations localised outlying $F_{ST}$ windows to the sweep region, whereas regular $F_{ST}$ showed outlying windows across the 10Mb simulation region.

Given the promising power of TwigScan on a simple simulated demographic history, we further extended our simulation study to include more variable selection coefficients and to approximate dog and wolf demographic history. We simulated two populations that split 10,000 generations ago: one with an effective population size of 10,000 (broadly representing the dog lineage) and one with an effective population size of 50,000 (broadly representing the wolf lineage). To do so, we used SLiM to simulate 10Mb genomes with a recombination rate of 5e-9 events per generation and no mutations, leaving it to run for 10,000 generations. We explored four different scenarios of selection each with four different selection coefficients (5%, 1%, 0.5% and 0.1%). These are shown in SI Figures 9 and 10.

In the first scenario, we inserted a mutation at the centre of the simulated region in the dog population 8,000 generations ago (corresponding approximately to 24,000 years). If this mutation reached 80% frequency, we changed the selection coefficient to 0 (labelled "8kgen"). In the second scenario, the mutation is maintained under selection until the completion of the simulation run (labelled "8kgen_sc"). The third and fourth scenarios are identical to the first and second, differing only in the timing of the selection: here, selection is initiated 3,000 generations before present (approx. 9,000 years ago) with either the selection coefficient changing to 0 once 80% frequency is reached (labelled "3kgen") or maintained until the completion of the simulation run (labelled "3kgen_sc").

In all cases, we then used pyslim (Gopalan et al. 2025) to recapitate this simulation. We chose an ancestral population size of 50,000 and a recombination rate of 5e-9. We then randomly sampled 100 diploid individuals from each population and added mutations with a rate of 1.25e-8 events per base per generation. We outputted the true tree sequence, as well as a VCF of the 400 sampled genomes.

Given these data, we inferred Relate trees with a constant haploid population size of 20,000 and mutation rate of 1.25e-8. We also converted the true tree sequences to Relate format using the relate_lib package. Finally, we then used TwigScan to compute $F_{ST}$ between the two populations using the function `Fst_blocks_from_Relate` for inferred ("Relate Trees") and true trees ("True Trees"). We used a generation cut-off of 14,000 and bin sizes of 10,000 or 50,000 bases (SI Figures 9 and 10), and compared to calculations only using mutations ("$F_{ST}$").

We show that for selection as old as 3k generations ago (9k years for dogs), calculating $F_{ST}$ using TwigScan provides a significant power improvement. For example, TwigScan calculated in 50kbp windows and for selection acting from 3k generations ago to present with a selection coefficient of 1%, we calculate a median z-score of 6.30 ([3.93,9.22] 95% CI) and 4.16 ([2.63,5.73] 95% CI) when applying TwigScan to true and Relate-inferred trees, respectively, compared to a z-score of 3.20 ([1.27,4.68] 95% CI) with traditional $F_{ST}$. Appreciable power remains with selection coefficients as low as 0.5%; we calculate a median z-score of 5.62 ([3.03,9.17] 95% CI) and 3.50 ([1.83,6.54] 95% CI) when applying TwigScan to true and Relate-inferred trees, respectively, compared to a z-score of 2.20 ([0.25,5.93] 95% CI) with traditional $F_{ST}$.

This diminishes somewhat for older selection, but was still marginally observed for selection 8k generations (24k years) ago till present; at a selection coefficient of 1%, we observe a median z-score of 3.87 ([1.40,5.29] 95% CI) and 3.58 ([1.38,4.82] 95% CI) when applying TwigScan to true and Relate-inferred trees, respectively, compared to a z-score of 3.20 with standard $F_{ST}$ ([1.15,4.78] 95% CI; SI Figures 10). We note that it is expected for TwigScan to be less effective for older selection occurring on the order of

Ne generations ago, given that many lineages will have coalesced at this point back in time.

In addition, to allow us to establish a conservative TwigScan $F_{ST}$ z-score cutoff signifying outlying windows putatively under selection, we simulate the dog and wolf genomes under neutrality (SI Figure 11). We simulated 38 wolf and dog chromosomes under the same demographic history as given above and using the inferred recombination maps from (Auton et al. 2013). We then recapitated the simulations, inferred trees using Relate, and additionally converted the true tree sequences to Relate format and calculated TwigScan $F_{ST}$ (using bin sizes of 50,000 bases and TwigScan cutoff of 14,000 generations) as described above. This showed that a z-score of five is only achieved for 0.07% of the windows, and a z-score of six is only achieved once (maximum z-score is 6.24) genome-wide (out of 46,358 windows in total) when using Relate trees. Further, the median z-score when applying TwigScan to Relate-inferred trees under neutrality is 0 ([-1.64,2.70] 95% CI).

**RST-Over Epochs Simulation**

When selection swiftly follows the birth of a mutation, the evidence for positive selection over the lifetime of the mutation is expected to be strong. Conversely, as the time between the onset of positive selection and the birth of the mutation increases, we expect that the evidence for positive selection over the lifetime of the mutation decreases. Here, we simulated positive selection with a varying time difference between the birth of a mutation and onset of selection. We then tested the Relate selection test over epochs (RST over epochs) on these simulations.

We used *SLiM* to simulate a population with constant diploid population size of 10,000 for 50,000 generations. We simulated 10Mb genomes with a recombination rate of 5e-9 events per base per generation and no mutations. We then recapitated the tree using *pyslim* and threw random mutations on the tree using a mutation rate of 1e-8.

We next read in the tree sequence file into *SLiM* once more and ran the simulation for another 1000 generations before initiating a selection event on a randomly chosen single mutation segregating at a frequency between 0.1-0.15 at generation 51,000. We ran this simulation 500 times for each simulated selection coefficient (0.5%, 1% and 5%). If the mutation was lost, we restarted the simulation with a new seed at generation 50,000. Once the selected mutation reached 80%, we terminated the simulation and outputted the true tree sequence and a VCF of 150 sampled genomes.

Given these data, we inferred Relate trees with a constant haploid population size of 20,000 and mutation rate of 1e-8. We also converted the true tree sequence to Relate format using the relate_lib package. Using RelateSelection, we inferred the evidence for positive selection over the lifetime of the mutation (when_mutation_has_freq2) and at time points following the distribution (0,10^seq(3,7,0.01)).

As the difference between the age of the mutation and the onset of selection, *i.e.*, the generation gap between mutation birth and selection increases, the evidence of selection as inferred over the lifetime of the mutation decreases relative to the evidence of selection inferred from discrete epochs surrounding the onset of positive selection (SI Figure 15). This was most obvious for the case of strong selection (selection coefficient simulated at 5%): there is appreciably higher evidence of positive selection at recent epochs than calculated over the lifetime of a mutation when there is as little as ~200-generation difference between selection onset and mutation birth.

### TwigScan applied to real data

##### ***TwigScan* applied to the 1000 Genomes Project data**

We applied the *TwigScan* function to the human Relate genealogies using phased genotypes from the combined 1000 Genomes Project and HGDP datasets aligned to GRCh38 (Koenig et al. 2024) (see section Relate genealogies). We used the *TwigScan* function implemented in the Twigstats R package (https://leospeidel.github.io/twigstats/reference/TwigScan.html) to compute $F_{ST}$ between the GBR and YRI with a window size of 50kb, using a cutoff time of 2000 generations (approx. 56k years). We compared these *TwigScan* $F_{ST}$ estimates with regular $F_{ST}$ estimates, computed also using the *TwigScan* function, but now setting options use_muts to TRUE, and specifying no time cutoff (t=Inf).

We show the result in Figure 3c and highlight two well-known loci under positive selection, *LCT* (chr2:135,787,850-135,837,184) and *SLC24A5* (chr15:48,120,990-48,142,672). We further list outlying regions with a TwigScan z-score greater than six in Supplementary Table 4, comparing these with previous selection scans.

##### ***TwigScan* applied to the Dog10K and 722g data**

Following a similar procedure, we applied the *TwigScan* function to the genome-wide genealogies inferred by Relate for the 722g and Dog10K datasets. We computed *TwigScan* $F_{ST}$ between the dogs and grey wolves using a window size of 50kb. We set the cutoff time to 14,000 generations (approx 42k years), chosen to predate the dog-grey wolf split. We again compared to regular $F_{ST}$ estimates by setting use_muts to TRUE and the time cutoff to infinity.

In Figure 4a, we highlighted outlying regions in the 722g genealogies, defined as a TwigScan z-score greater than six. This cutoff was chosen based on our earlier neutral simulations, indicating that a TwigScan z-score greater than six was very unlikely under neutrality. A list of these outlying regions putatively under selection in dogs are listed in Supplementary Table 5. In this table we list genes located inside the selection-scans windows as well as within 250k base-pair of the focal window. These neighbouring windows are also highlighted in the Manhattan plot in Fig 4a.

To compare to the Dog10K dataset, we additionally computed an empirical p-value within the Dog10K dataset, by first lifting over these outlying regions using the UCSC liftover tool (https://genome.ucsc.edu/cgi-bin/hgLiftOver) and then computing how many genome-wide windows in Dog10K had at least the observed TwigScan $F_{ST}$ score. These empirical p-values are also listed in Supplementary Table 5, alongside an empirical p-value where we used the minimum across at most two windows upstream or downstream. We chose this approach because the window definition is approximate, and liftover coordinates may therefore be offset by some distance. All empirical p-values were <0.05, with *AMY2B*/*RNPC3* achieving <1e-4. When we allowed two adjacent windows, the empirical p-value was 2.28E-05, which equalled the top window genome-wide (total number of windows 43,808). The locus including *MED13L* had an empirical p-value of 0.007, and if we allowed two adjacent windows 5.48E-04.

### Detailed analysis of the *AMY2B/RNPC3* locus

##### Dataset

We carried out the following analyses using the 722g extended dataset as all the additional ancient and modern canids were aligned to canFam3.1.

##### Linkage disequilibrium around the *AMY2B*/*RNPC3* locus

To better characterise the haplotype present in most dogs at *RNPC3*, we identified a block of SNPs in perfect LD as follows. Using the phased genotype data for the 722g dataset, we chose an initial SNP with the greatest difference in allele frequency between modern dogs and modern wolves within the *RNPC3* gene (chr6: 46960033-47044272). We next computed LD (measured as $r^2$) between this SNP and every other SNP in a 200kbp region centered on it. We then selected the top 50 SNPs with the highest $r^2$ score and computed an artificial pseudo-SNP by summing genotype status across the 50 SNPs with highest LD. We iterated this procedure, computing LD between this artificial SNP and each SNP within the 200kbp region and then updating the artificial SNP. We repeated this scheme until convergence. Once converged, we again computed LD to all mutations within the 200kb window and obtained 305 SNPs in perfect LD (r2 = 1) to each other (in modern dogs and wolves) tagging the haplotype putatively under selection.

##### Visualisation of the RNPC3 haplotype across wolf-like canids

We visualised the *RNPC3* SNPs in perfect LD in a genotype matrix across canids, including dogs, grey wolves, coyotes, red wolves, African golden wolves, golden jackals, dholes, Ethiopian wolves, and African wild dogs (Liu et al. 2018; Gopalakrishnan et al.

2018; Plassais et al. 2019; Kardos et al. 2018; Sinding et al. 2018). We polarised alleles relative to a high-coverage northwestern Chinese wolf (WolfTibetan04). We separated Arctic dog breeds and dingoes from other dogs, as these breeds carry the focal haplotype at low frequency. To minimise missingness in the genotype matrix, we excluded samples with over 5% missingness in the dog and grey wolf groups as they each contain a relatively large number of samples, before filtering out any SNPs with over 50% missingness in African golden wolves. This resulted in 311 SNPs that were plotted in a genotype matrix (Fig 4f).

**Cross-species relationships at *RNPC3* and *EPAS1***

To assess cross-canid divergence relationships, we computed pairwise $f_2$-statistics between extant wolf-like canids in windows across the genome. We divided the genome into exact non-overlapping 50kb regions. Additionally, we extracted the focal regions for *RNPC3* and *EPAS1*, centred on chr6:47,000,000 and chr10:48,600,000 in CanFam3.1 (SI Figure 13). We then computed a single $f_2$-statistic (computed here simply as the mean squared difference of allele frequencies) for each window between all pairs of species. We used the genome-wide windows as a null to the $f_2$ value in the *RNPC3* and *EPAS1* windows of interest. For each focal window of interest, we excluded 10Mb upstream and downstream from the null distribution.

We show the results in SI Figure 13. At *RNPC3*, dogs and African golden wolves do indeed carry a deeply divergent haplotype that appears outlying even when compared to more distant wolf-like canids such as dholes and African wild dogs. At *EPAS1*, Tibetan dogs and wolves carry a deeply divergent haplotype that is slightly less diverged, appearing outlying compared to Ethiopian wolves and African wild dogs but not dholes. In addition, we find that pairwise relationships between other canids not impacted by introgression (i.e. not dogs or African golden wolves for RNPC3, not Tibetan dogs or wolves for EPAS1) are not unusually diverged to each other, suggesting that the *RNPC3* and *EPAS1* windows are not outlying with respect to local mutation rates or other properties that could increase local $f_2$-statistics across species.

For *EPAS1*, we chose individuals Wolf42, WolfTibetan01, WolfTibetan02, WolfTibetan03, WolfTibetan04 and individuals ChineseIndigenousDog02, ChineseIndigenousDog04, VillDog_China30, VillDog_China41, ChineseIndigenousDog11, VillDog_China46, VillDog_China31, TibetanMastiff11, TibetanMastiff10, TibetanMastiff08, VillDog_China40, VillDog_China44, VillDog_China32, TibetanMastiff02.

**Copy number variation at *AMY2B***

To understand the relationship between the introgressed haplotype at *RNPC3* and *AMY2B* copy number status, we computed copy number variation at *AMY2B* in present-day and ancient canids. We aligned against a high-quality grey wolf assembly (Sinding et al. 2021) using bwa mem. We then computed per-base coverage around *AMY2B* using samtools for the region HG994389.1:45250000-50250000. We computed coverage in the interval [47745000, 47755000] containing the *AMY2B* gene using the command samtools depth -aa -Q 20. We then also computed the coverage in flanking regions after excluding 100kb around 47,750,000 using the same command. We divided these two estimates to obtain a relative coverage at the *AMY2B* gene. We quantified uncertainty using a block-bootstrap with block size of 500bp, where we resampled blocks from the focal and flanking regions 1000 times and computed the 2.5th and 97.5th percentiles of the relative coverage at *AMY2B*.

**Allele frequency trajectory at the *RNPC3* locus**

Most ancient individuals are only sequenced to low coverage and are called as pseudo-haploid. We therefore called diploid haplotype carrier status by computing the mean genotype across SNPs in perfect LD within the *RNPC3* haplotype after polarising those genotypes to the introgressed haplotype, multiplied by 2 and rounded to the nearest integer, to obtain a diploid call (Supplementary Table 6). The majority of our samples were from Western Eurasia and therefore we restricted the samples to this region.

We used these genotypes alongside sampling ages of ancient dogs to estimate allele frequency trajectories and selection coefficients using CLUES2. We used diploid calls for every ancient individual as input and converted sampling ages from years to generations assuming a generation time of 3 years in dogs. We used the --ancientSamps argument in CLUES2 to parse the genotype data.

The present-day derived allele frequency was estimated from the modern samples and used as input (popFreq=0.95). Analyses were run with a maximum time cutoff of 5,500 generations (~16kya) corresponding to the time span of our aDNA time series. We assumed a model of constant selection, such that a single selection coefficient was estimated over the entire time interval. We used coalescence rates inferred for dogs from the 722g genealogies. We used a discretisation parameter of f = 750, all remaining parameters were left at their default values.

Genomic History Reveals a Dual Ancestry of Dogs.” *Nature* 607 (7918): 313–20.

Botigué, Laura R., Shiya Song, Amelie Scheu, et al. 2017. “Ancient European Dog Genomes Reveal Continuity since the Early Neolithic.” *Nature Communications* 8 (1): 16082.

Carter, Evelyn J., Eirlys E. Tysall, Jason A. Hodgson, and Andrea Manica. 2026. “*Tidypopgen* : Tidy Population Genetics in R.” *Methods in Ecology and Evolution* 17 (2): 480–87.

Chavez, Daniel E., Ilan Gronau, Taylor Hains, et al. 2022. “Comparative Genomics Uncovers the Evolutionary History, Demography, and Molecular Adaptations of South American Canids.” *Proceedings of the National Academy of Sciences of the United States of America* 119 (34): e2205986119.

Chen, Shifu, Yanqing Zhou, Yaru Chen, and Jia Gu. 2018. “Fastp: An Ultra-Fast All-in-One FASTQ Preprocessor.” *Bioinformatics (Oxford, England)* 34 (17): i884–90.

Chen, Siwei, Laurent C. Francioli, Julia K. Goodrich, et al. 2024. “A Genomic Mutational Constraint Map Using Variation in 76,156 Human Genomes.” *Nature* 625 (7993): 92–100.

Danecek, Petr, James K. Bonfield, Jennifer Liddle, et al. 2021. “Twelve Years of SAMtools and BCFtools.” *GigaScience* 10 (2): giab008.

Delaneau, Olivier, Jean François Zagury, Matthew R. Robinson, Jonathan L. Marchini, and Emmanouil T. Dermitzakis. 2019. “Accurate, Scalable and Integrative Haplotype Estimation.” *Nature Communications* 10 (1): 24–29.

Fan, Zhenxin, Pedro Silva, Ilan Gronau, et al. 2016. “Worldwide Patterns of Genomic Variation and Admixture in Gray Wolves.” *Genome Research* 26 (2): 163–73.

Fellows Yates, James A., Thiseas C. Lamnidis, Maxime Borry, et al. 2021. “Reproducible, Portable, and Efficient Ancient Genome Reconstruction with Nf-Core/Eager.” *PeerJ* 9 (e10947): e10947.

Feuerborn, Tatiana R., Alberto Carmagnini, Robert J. Losey, et al. 2021. “Modern Siberian Dog Ancestry Was Shaped by Several Thousand Years of Eurasian-Wide Trade and Human Dispersal.” *Proceedings of the National Academy of Sciences of the United States of America* 118 (39): e2100338118.

Frantz, Laurent A. F., Victoria E. Mullin, Maud Pionnier-Capitan, et al. 2016. “Genomic and Archaeological Evidence Suggest a Dual Origin of Domestic Dogs.” *Science (New York, N.Y.)* 352 (6290): 1228–31.

Freedman, Adam H., Ilan Gronau, Rena M. Schweizer, et al. 2014. “Genome Sequencing Highlights the Dynamic Early History of Dogs.” *PLoS Genetics* 10 (1): e1004016.

Girdland-Flink, Linus, Anders Bergström, Jan Storå, et al. 2025. “Gray Wolves in an Anthropogenic Context on a Small Island in Prehistoric Scandinavia.” *Proceedings of the National Academy of Sciences of the United States of America* 122 (48): e2421759122.

Gopalakrishnan, Shyam, Mikkel-Holger S. Sinding, Jazmín Ramos-Madrigal, et al. 2018. “Interspecific Gene Flow Shaped the Evolution of the Genus Canis.” *Current Biology: CB* 28 (21): 3441-3449.e5.

Gopalan, Shyamalika, Murillo F. Rodrigues, Peter L. Ralph, and Benjamin C. Haller. 2025. “Bridging Forward-in-Time and Coalescent Simulations Using Pyslim.” In *Evolutionary Biology*, Biorxiv;2025.09.30.679676v1. bioRxiv, October 1. https://www.biorxiv.org/content/10.1101/2025.09.30.679676v1.

Griffiths, R. C., and Simon Tavaré. 1998. “The Age of a Mutation in a General Coalescent Tree.” *Communications in Statistics* 14 (1–2): 273–95.

Kardos, Marty, Mikael Åkesson, Toby Fountain, et al. 2018. “Genomic Consequences of Intensive Inbreeding in an Isolated Wolf Population.” *Nature Ecology & Evolution* 2 (1): 124–31.

Kelleher, Jerome, Alison M. Etheridge, and Gilean McVean. 2016. “Efficient Coalescent Simulation and Genealogical Analysis for Large Sample Sizes.” *PLoS Computational Biology* 12 (5): e1004842.

Koenig, Zan, Mary T. Yohannes, Lethukuthula L. Nkambule, et al. 2024. “A Harmonized Public Resource of Deeply Sequenced Diverse Human Genomes.” *Genome Research* 34 (5): 796–809.

Li, Heng. 2013. “Aligning Sequence Reads, Clone Sequences and Assembly Contigs with BWA-MEM.” In *arXiv [q-Bio.GN]*. March 16. arXiv. https://doi.org/10.48550/arXiv.1303.3997.

Li, Heng, and Richard Durbin. 2009. “Fast and Accurate Short Read Alignment with Burrows-Wheeler Transform.” *Bioinformatics* 25 (14): 1754–60.

Lin, Audrey T., Liz Hammond-Kaarremaa, Hsiao-Lei Liu, et al. 2023. “The History of Coast Salish ‘Woolly Dogs’ Revealed by Ancient Genomics and Indigenous Knowledge.” *Science (New York, N.Y.)* 382 (6676): 1303–8.

Liu, Yan-Hu, Lu Wang, Tao Xu, et al. 2018. “Whole-Genome Sequencing of African Dogs Provides Insights into Adaptations against Tropical Parasites.” *Molecular Biology and Evolution* 35 (2): 287–98.

Lu, Dongsheng, Haiyi Lou, Kai Yuan, et al. 2016. “Ancestral Origins and Genetic History of Tibetan Highlanders.” *American Journal of Human Genetics* 99 (3): 580–94.

Mak, Sarah Siu Tze, Shyam Gopalakrishnan, Christian Carøe, et al. 2017. “Comparative Performance of the BGISEQ-500 vs Illumina HiSeq2500 Sequencing

Platforms for Palaeogenomic Sequencing.” *GigaScience* 6 (8): 1–13.

Marco-Sola, Santiago, and Paolo Ribeca. 2015. “Efficient Alignment of Illumina-like High-Throughput Sequencing Reads with the GEnomic Multi-Tool (GEM) Mapper.” *Current Protocols in Bioinformatics* 50 (1): 11.13.1-11.13.20.

Marsh, William A., Lachie Scarsbrook, Eren Yüncü, et al. 2026. “Dogs Were Widely Distributed across Western Eurasia during the Palaeolithic.” *Nature* 651 (8107): 995–1003.

Meadows, Jennifer R. S., Jeffrey M. Kidd, Guo-Dong Wang, et al. 2023. “Genome Sequencing of 2000 Canids by the Dog10K Consortium Advances the Understanding of Demography, Genome Function and Architecture.” *Genome Biology* 24 (1): 187.

Meyer, Matthias, Martin Kircher, Marie-Theres Gansauge, et al. 2012. “A High-Coverage Genome Sequence from an Archaic Denisovan Individual.” *Science* 338 (6104): 222–26.

Neukamm, Judith, Alexander Peltzer, and Kay Nieselt. 2021. “DamageProfiler: Fast Damage Pattern Calculation for Ancient DNA.” *Bioinformatics (Oxford, England)* 37 (20): 3652–53.

Ní Leathlobhair, Máire, Angela R. Perri, Evan K. Irving-Pease, et al. 2018. “The Evolutionary History of Dogs in the Americas.” *Science (New York, N.Y.)* 361 (6397): 81–85.

Niemann, Jonas, Shyam Gopalakrishnan, Nobuyuki Yamaguchi, et al. 2021. “Extended Survival of Pleistocene Siberian Wolves into the Early 20th Century on the Island of Honshū.” *iScience* 24 (1): 101904.

Ostrander, Elaine A., Guo-Dong Wang, Greger Larson, et al. 2019. “Dog10K: An International Sequencing Effort to Advance Studies of Canine Domestication, Phenotypes and Health.” *National Science Review* 6 (4): 810–24.

Patterson, Nick, Priya Moorjani, Yontao Luo, et al. 2012. “Ancient Admixture in Human History.” *Genetics* 192 (3): 1065–93.

Peltzer, Alexander, Günter Jäger, Alexander Herbig, et al. 2016. “EAGER: Efficient Ancient Genome Reconstruction.” *Genome Biology* 17 (1): 60.

Perri, Angela R., Kieren J. Mitchell, Alice Mouton, et al. 2021. “Dire Wolves Were the Last of an Ancient New World Canid Lineage.” *Nature* 591 (7848): 87–91.

“Picard.” n.d. Accessed August 12, 2026. https://broadinstitute.github.io/picard/.

Plassais, Jocelyn, Jaemin Kim, Brian W. Davis, et al. 2019. “Whole Genome Sequencing of Canids Reveals Genomic Regions under Selection and Variants Influencing Morphology.” *Nature Communications* 10 (1): 1489.

Prüfer, Kay, Fernando Racimo, Nick Patterson, et al. 2014. “The Complete Genome Sequence of a Neanderthal from the Altai Mountains.” *Nature* 505 (7481): 43–49.

Ramos-Madrigal, Jazmín, Mikkel-Holger S. Sinding, Christian Carøe, et al. 2021. “Genomes of Pleistocene Siberian Wolves Uncover Multiple Extinct Wolf Lineages.” *Current Biology: CB* 31 (1): 198-206.e8.

Sabeti, Pardis C., Patrick Varilly, Ben Fry, et al. 2007. “Genome-Wide Detection and Characterization of Positive Selection in Human Populations.” *Nature* 449 (7164): 913–18.

Schubert, Mikkel, Stinus Lindgreen, and Ludovic Orlando. 2016. “AdapterRemoval v2: Rapid Adapter Trimming, Identification, and Read Merging.” *BMC Research Notes* 9 (1): 88.

Segawa, Takahiro, Takahiro Yonezawa, Hiroshi Mori, et al. 2022. “Paleogenomics Reveals Independent and Hybrid Origins of Two Morphologically Distinct Wolf Lineages Endemic to Japan.” *Current Biology: CB* 32 (11): 2494-2504.e5.

Sinding, Mikkel-Holger S., Shyam Gopalakrishan, Filipe G. Vieira, et al. 2018. “Population Genomics of Grey Wolves and Wolf-like Canids in North America.” *PLoS Genetics* 14 (11): e1007745.

Sinding, Mikkel-Holger S., Shyam Gopalakrishnan, Jazmín Ramos-Madrigal, et al. 2020. “Arctic-Adapted Dogs Emerged at the Pleistocene-Holocene Transition.” *Science (New York, N.Y.)* 368 (6498): 1495–99.

Sinding, Mikkel-Holger S., Shyam Gopalakrishnan, Katrine Raundrup, et al. 2021. “The Genome Sequence of the Grey Wolf, Canis Lupus Linnaeus 1758.” *Wellcome Open Research* 6 (November): 310.

Skoglund, Pontus, Erik Ersmark, Eleftheria Palkopoulou, and Love Dalén. 2015. “Ancient Wolf Genome Reveals an Early Divergence of Domestic Dog Ancestors and Admixture into High-Latitude Breeds.” *Current Biology: CB* 25 (11): 1515–19.

Speidel, Leo, Marie Forest, Sinan Shi, and Simon R. Myers. 2019. “A Method for Genome-Wide Genealogy Estimation for Thousands of Samples.” *Nature Genetics* 51 (9): 1321–29.

St. John, John. n.d. *SeqPrep: Tool for Stripping Adaptors and/or Merging Paired Reads with Overlap into Single Reads*. Github. Accessed August 12, 2026. https://github.com/jstjohn/SeqPrep.

vonHoldt, Bridgett M., James A. Cahill, Zhenxin Fan, et al. 2016. “Whole-Genome Sequence Analysis Shows That Two Endemic Species of North American Wolf Are Admixtures of the Coyote and Gray Wolf.” *Science Advances* 2 (7): e1501714.

Wang, Guo-Dong, Weiwei Zhai, He-Chuan Yang, et al. 2016. “Out of Southern East Asia: The Natural History of Domestic Dogs across the World.” *Cell Research* 26 (1):

21–33.

Zhang, Shao-Jie, Jilong Ma, Meritxell Riera, et al. 2025. “Determinants of de Novo Mutations in Extended Pedigrees of 43 Dog Breeds.” *Genome Biology* 26 (1): 305.

### Supplementary Figures

#### SI Figure 1 Datasets

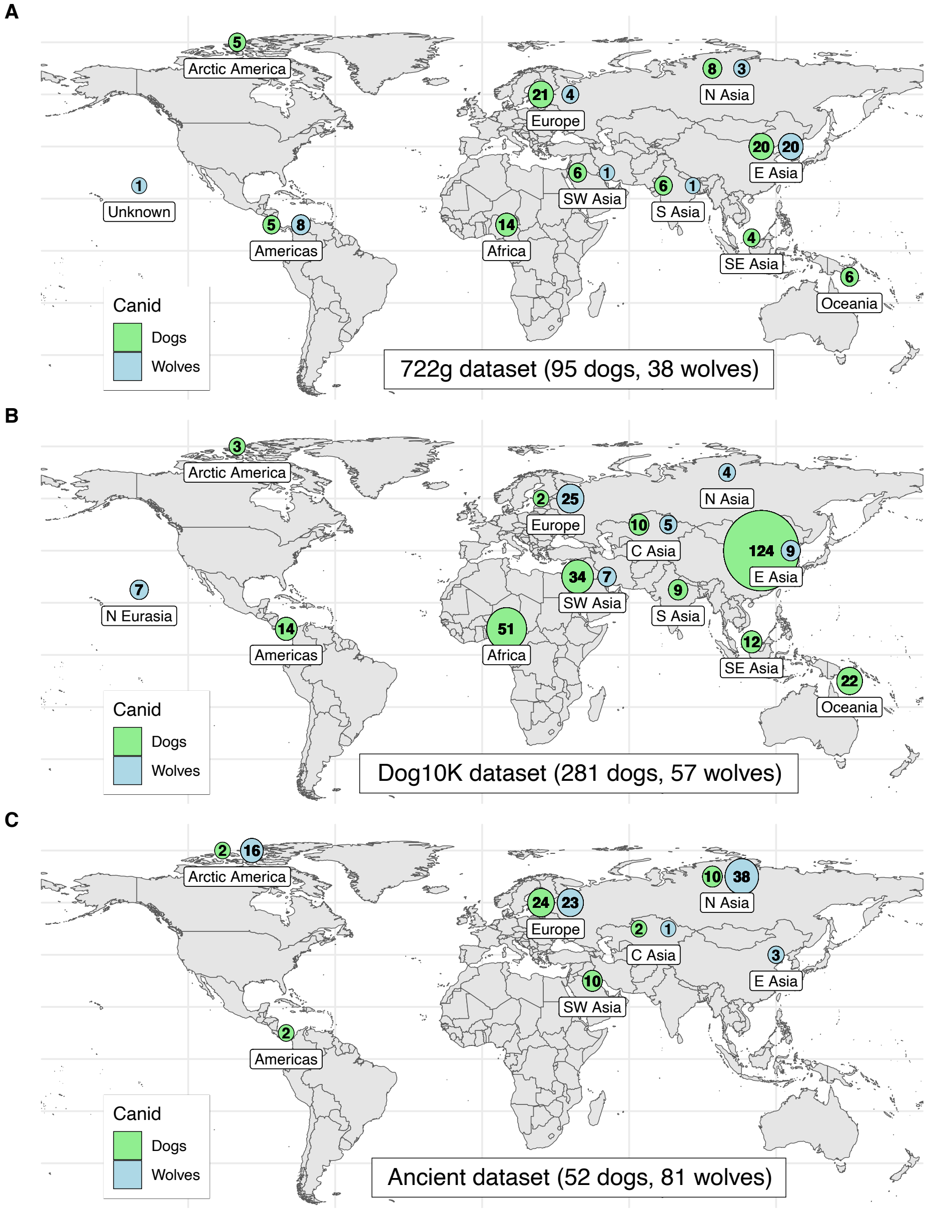


Geographic location of samples in the 722g (a) and Dog10K (b) datasets which contain present-day wolves and dogs, and a dataset of ancient dogs and wolves (c). In the Dog10K dataset, all samples of unknown origin are north Eurasian.

**SI Figure 2 PCA of modern dog samples mapped to canFam3.1**

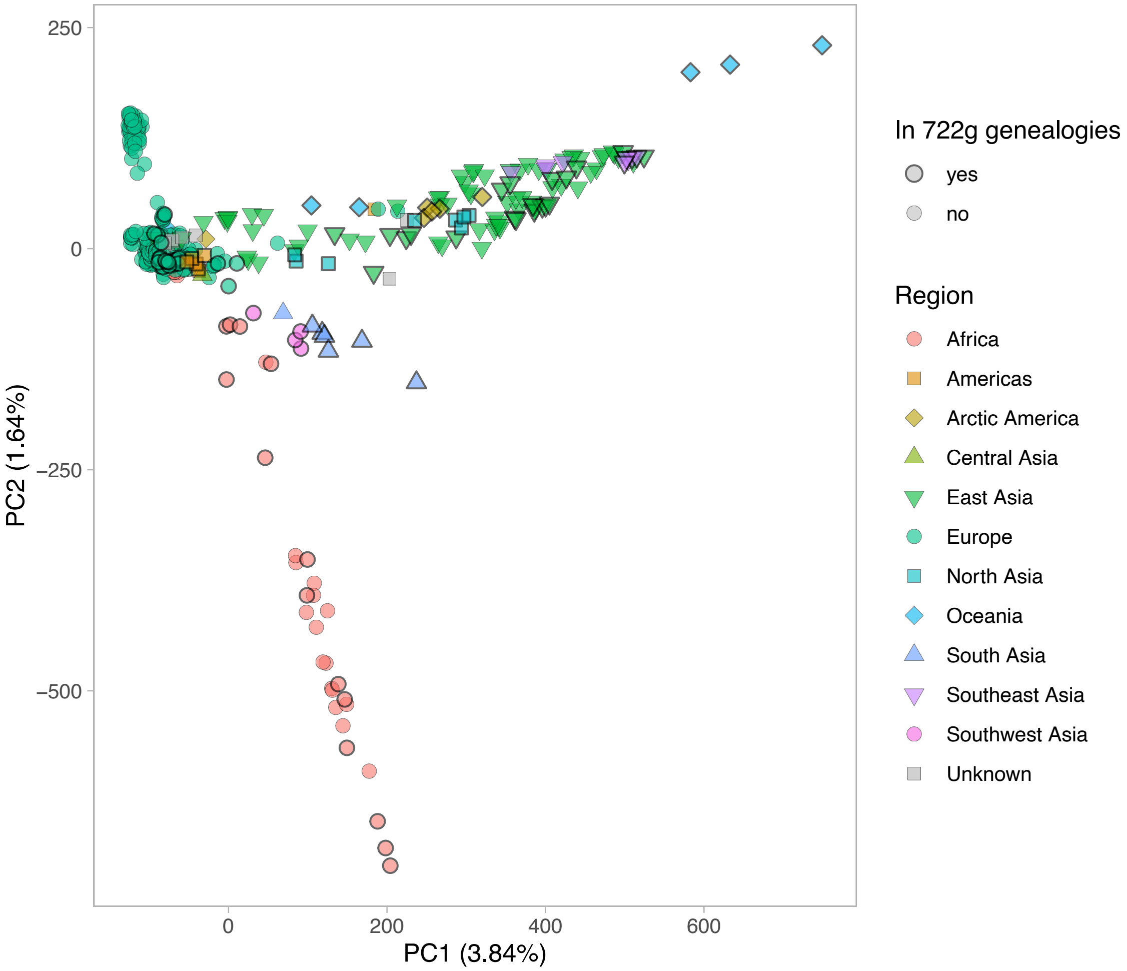


PCA for all dog samples aligned to canFam3.1 (722g and additional downloaded dog genomes, SI Table 3). The full dataset was filtered by removing individuals with less than 65% (45M) of genotyped SNPs, and SNPs were removed if they had over 10% missingness in the remaining individuals, resulting in 611,160 SNPs. Genotypes were then imputed and used to compute the PCA using *tidypopgen*. The subset of 722g samples used for genealogy inference are highlighted with black circles.

**SI Figure 3 Context-dependent mutation rates**

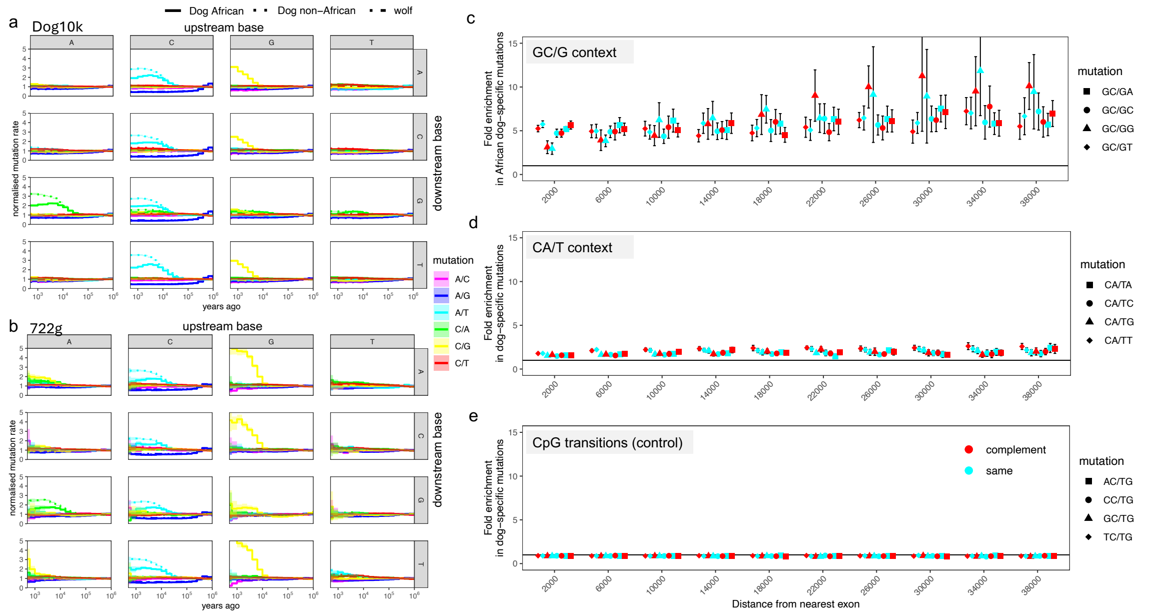


**a,** Mutation rates for 96 triplet mutations inferred using the Dog10K genealogies. Linetypes show mutation rates for grey wolves, African dogs, and non-African dogs. Column facets correspond to the upstream base, row facets to the downstream base. **b,** Same as a, but for the 722g genealogies. **c**, Fold enrichment of mutations unique to African dogs relative to mutations not unique to African dogs, in context CG/GG. We restricted to mutations that are younger than 30k years, and stratify by distance to the nearest exon (Supplementary Information). We computed this enrichment separately for 192 trinucleotide contexts and show the reverse complements in separate colours, to illustrate any strand effects. Solid black line indicates 1 (no enrichment). We show 2 standard errors. **d**, Same as c but for context CA/CT, computing enrichment of mutations unique to dogs. **e**, Same as c but for CpG contexts, where we do not expect an enrichment specific to dogs.

**SI Figure 4 Context-dependent mutation enrichment in each individual dog sample**

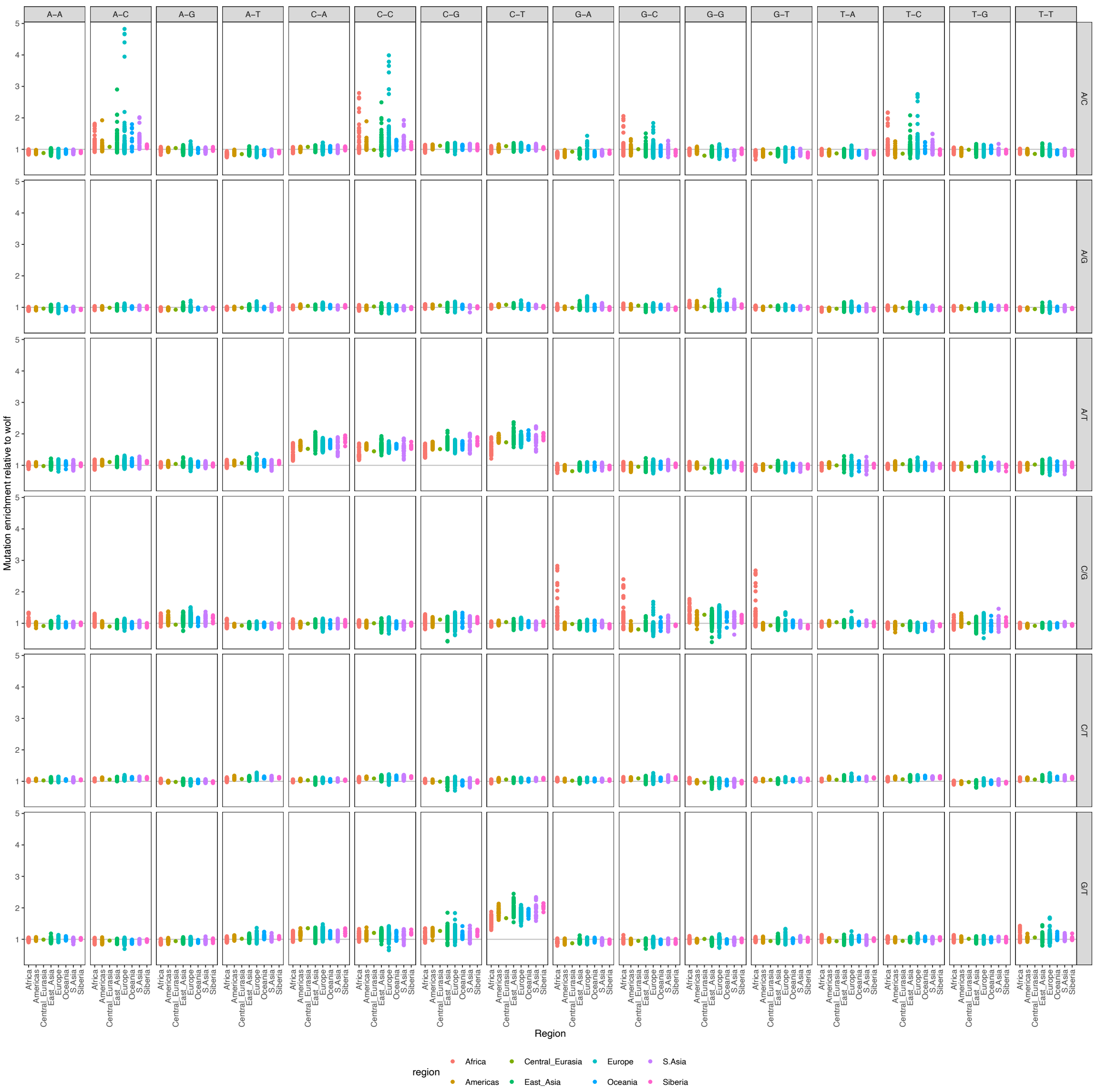


Mutation count enrichment in all 96 categories relative to grey wolves, computed from genotypes for every present-day dog sample in our dataset (Supplementary Information). We used SNPs carried in at least two individuals (as het or hom) to avoid sequencing artefacts. We computed enrichment by first counting the number of mutations in each trinucleotide context per dog sample. We then divided by the corresponding count computed across all grey wolves and normalised such that across the 96 categories, the median enrichment equals 1 for each sample. Column facets show upstream and downstream nucleotides, and row facets show mutation type. Colours correspond to geographic region assignment of each dog sample.

**SI Figure 5 GC-biased gene conversion**

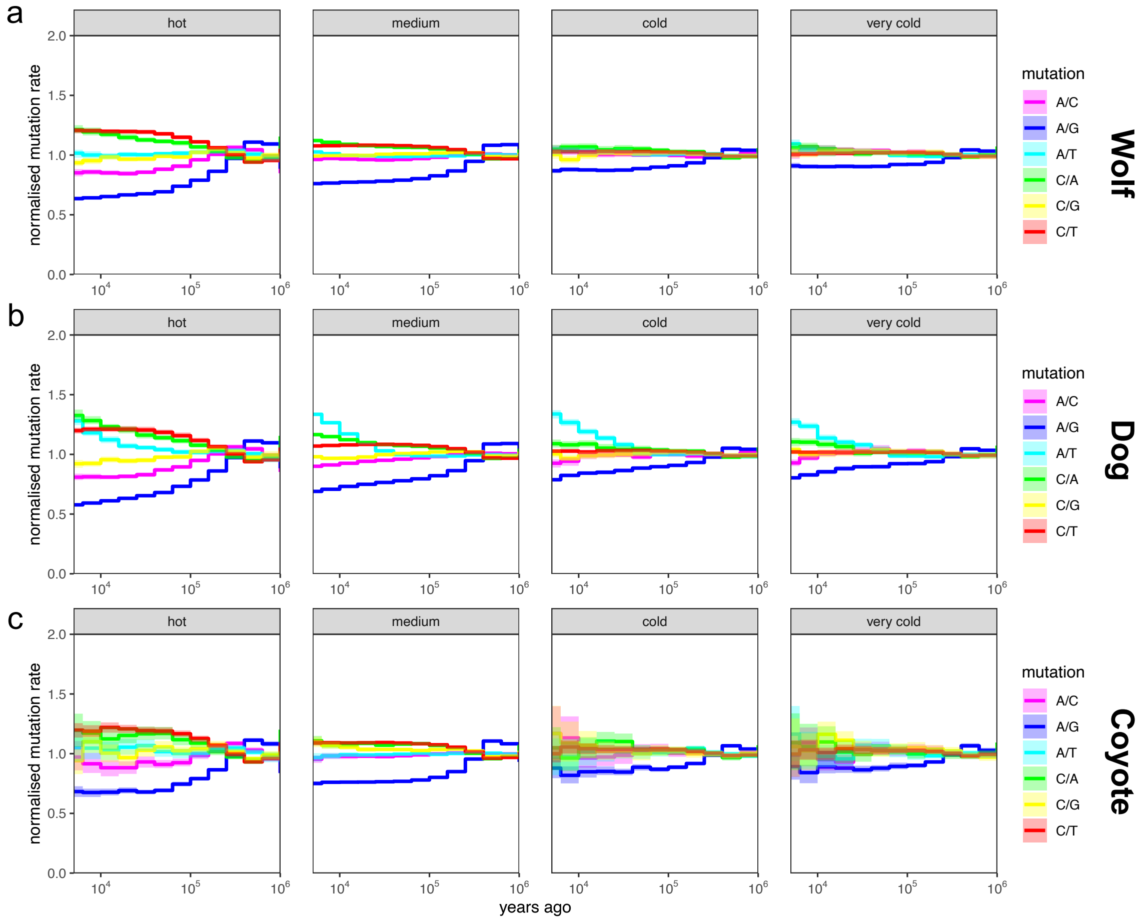


Evidence of GC biased gene conversion in **a,** grey wolves, **b,** dogs, and **c,** coyotes. We stratified the genome by recombination hotness using the recombination map inferred by Ref. (Auton et al. 2013) and computed each of the six single nucleotide mutation rates. We replaced any regions with an inferred recombination rate of 0 by the genome-wide average, as these are typically only indicating missing data. Very cold recombination regions correspond to the bottom 5%, cold corresponds to 5% - 10%, medium to 10% - 95%, and hot to the top 5%.

#### SI Figure 6 Comparison to de-novo mutations

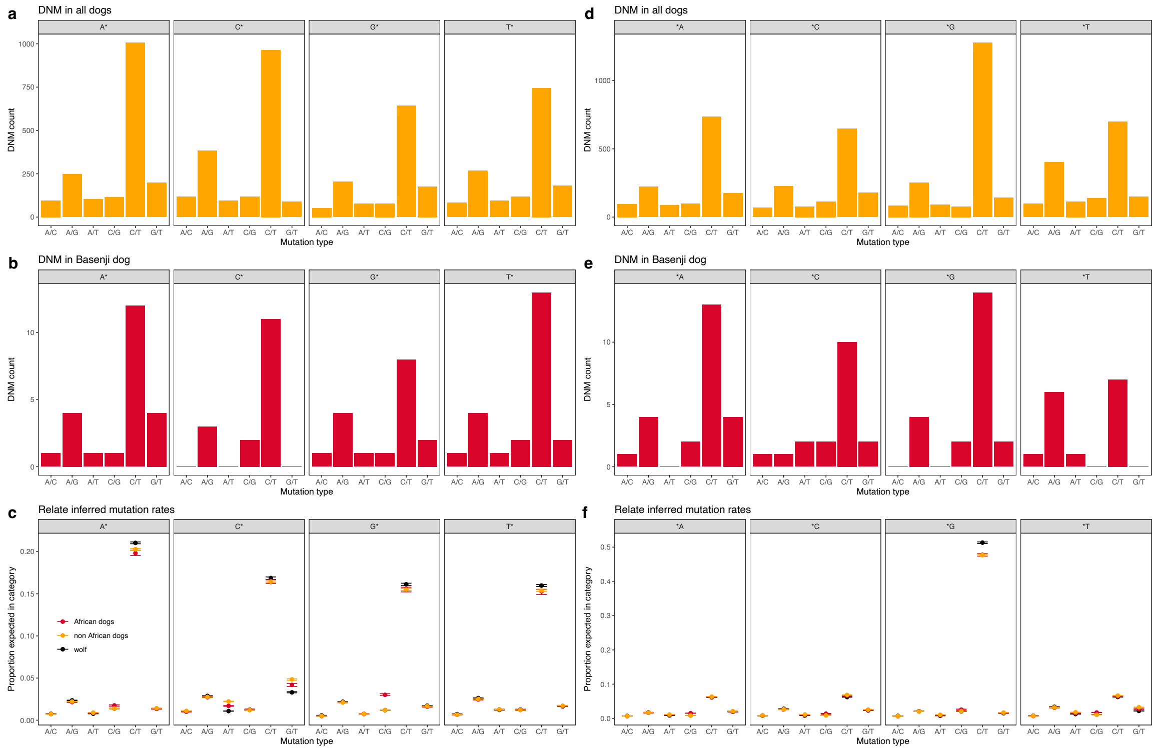


**a,** De-novo mutation counts across all dog breeds using data from Ref. (Zhang et al. 2025). We define mutation categories by upstream nucleotide and focal mutation type, accounting for strand symmetries. **b**, Same as a, but only for the Basenji dog breed. **c,** Mutation rates inferred by Relate. We focussed on the time period 1,000 - 10,000 years ago, and computed the proportion of mutations expected in each category, in grey wolves, non-African dogs, and African dogs (Supplementary Information). **d - f,** Same as **a - c,** but now depending on the downstream nucleotide.

**SI Figure 7 Species-wide TMRCAs**

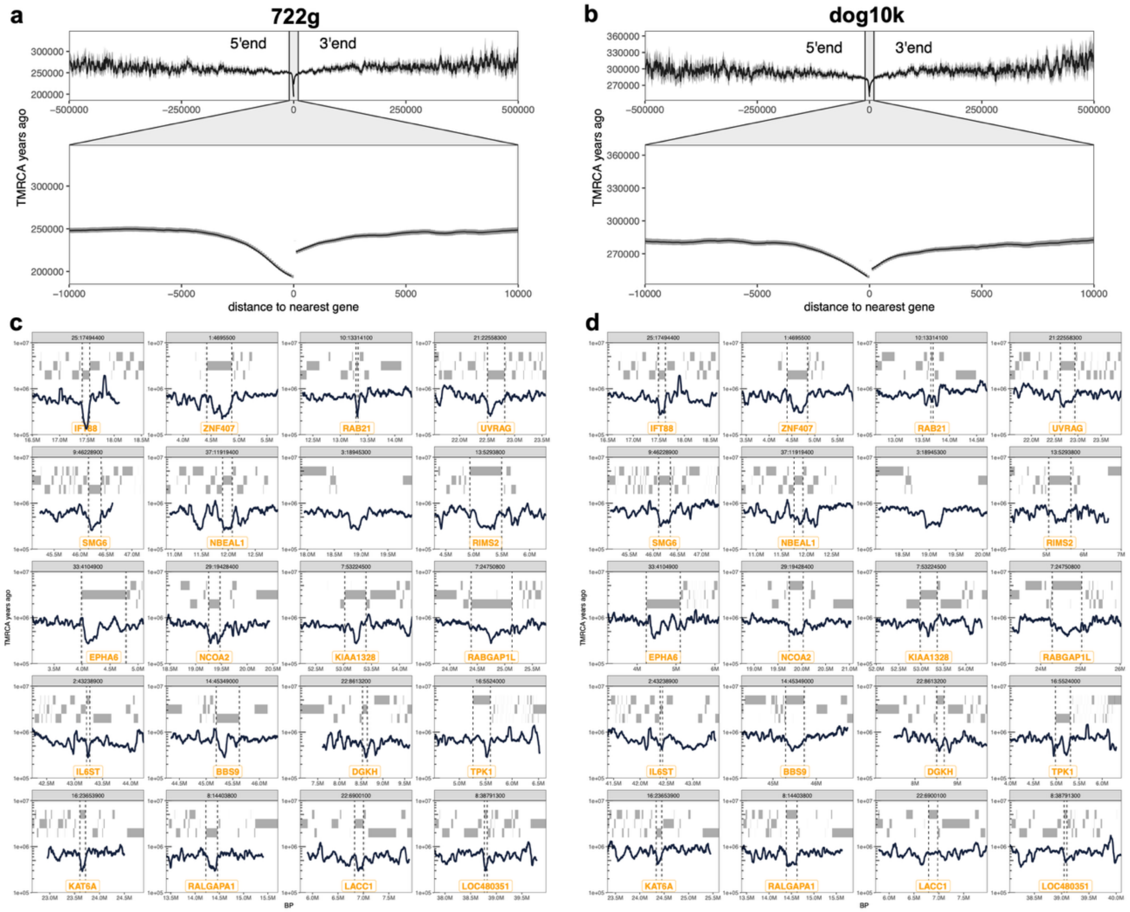


**a,** Average TMRCA of all grey wolves and dogs in the 722g genealogies by distance to the nearest gene. We computed distance to the 5' or 3' end of the gene and any bases falling inside a gene have distance zero. Error bars show 1.96 standard errors of the mean. **b,** Same as **a** but for the Dog10K genealogies. **c,** The top 20 regions with the youngest TMRCAs in the 722g genealogies (Supplementary Information). **d,** Same regions as in **c,** but now showing TMRCAs in the Dog10K genealogies. In **c** and **d**, grey rectangles correspond to protein coding genes. The minimum TMRCA overlaps a gene in all but one region and the names of these genes are shown in orange, with dashed grey lines delineating their borders.

**SI Figure 8 Comparison of TwigScan and XP-EHH**

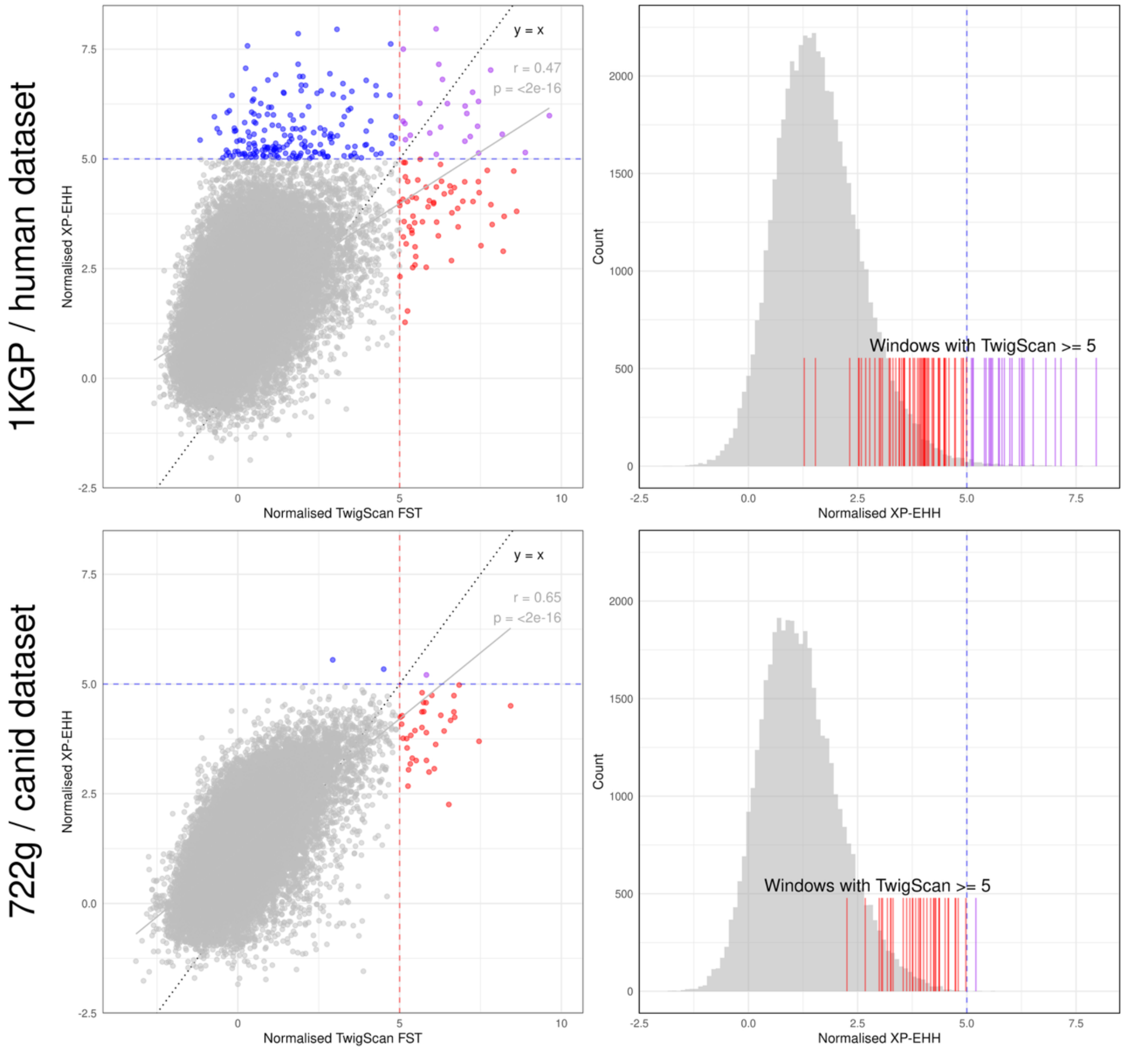


Comparison of TwigScan results with XP-EHH for humans and canids (Sabeti et al. 2007). The SNP-level XP-EHH results were split in 50kbp windows and the window value was determined by calculating the mean of the top five SNPs in the window. Outlier windows identified using TwigScan broadly lie in the tail of the XP-EHH distribution.

**SI Figure 9: TwigScan performance in simulated dog loci under selection (10kb windows).**

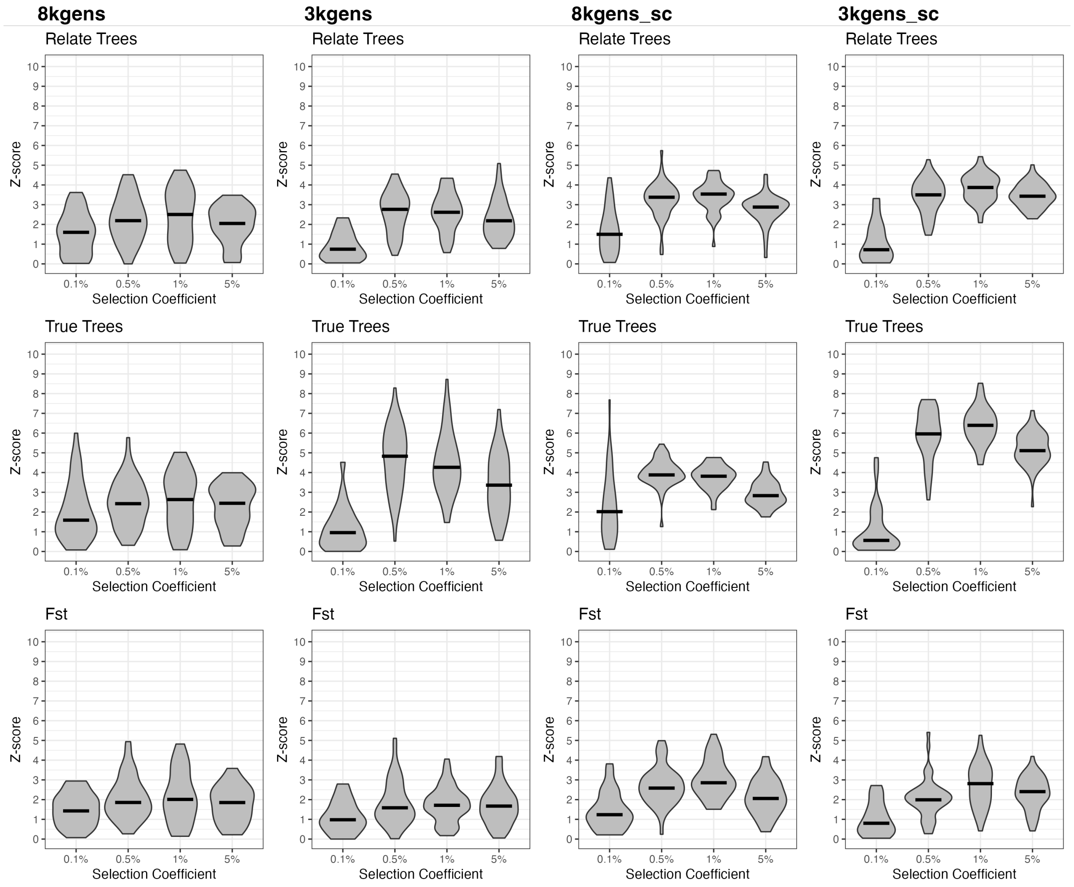


Distribution of z-scores (y-axis) calculated on 10,000 base pair windows surrounding the mutation under selection (centre of simulated genome segment) for TwigScan calculated on Relate-inferred trees (top panel), true tree sequences (middle panel), and Fst calculated using mutations (bottom panel). Shown for four simulated selection coefficients (x-axis) and four simulated selection scenarios (see Supplementary Information, section TwigScan simulations).

**SI Figure 10: TwigScan performance in simulated dog loci under selection (50kb windows).**

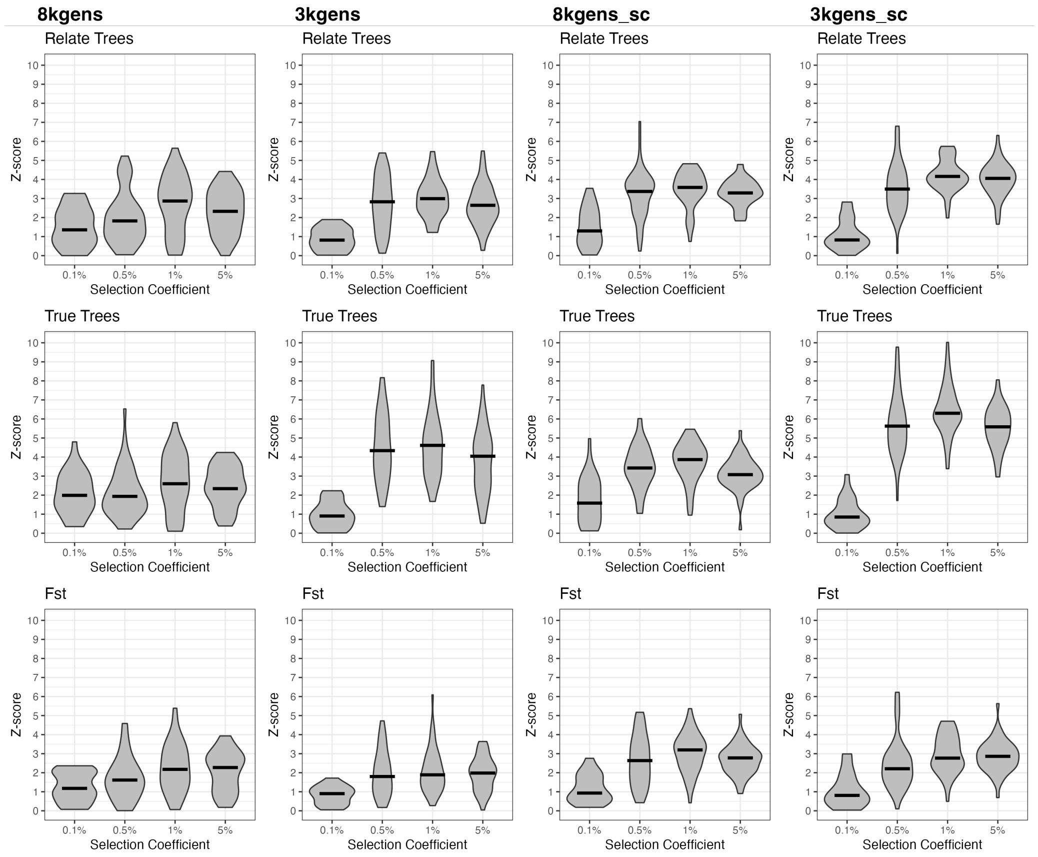


Distribution of z-scores (y-axis) calculated on 50,000 base pair windows surrounding the mutation under selection (centre of simulated genome segment) for TwigScan calculated on Relate-inferred trees (top panel), true tree sequences (middle panel), and Fst calculated using mutations (bottom panel). Shown for four simulated selection coefficients (x-axis) and four simulated selection scenarios (see Supplementary Information, section TwigScan simulations).

**SI Figure 11: Comparison of TwigScan $F_{ST}$ and standard $F_{ST}$ on neutrally simulated canid genomes**

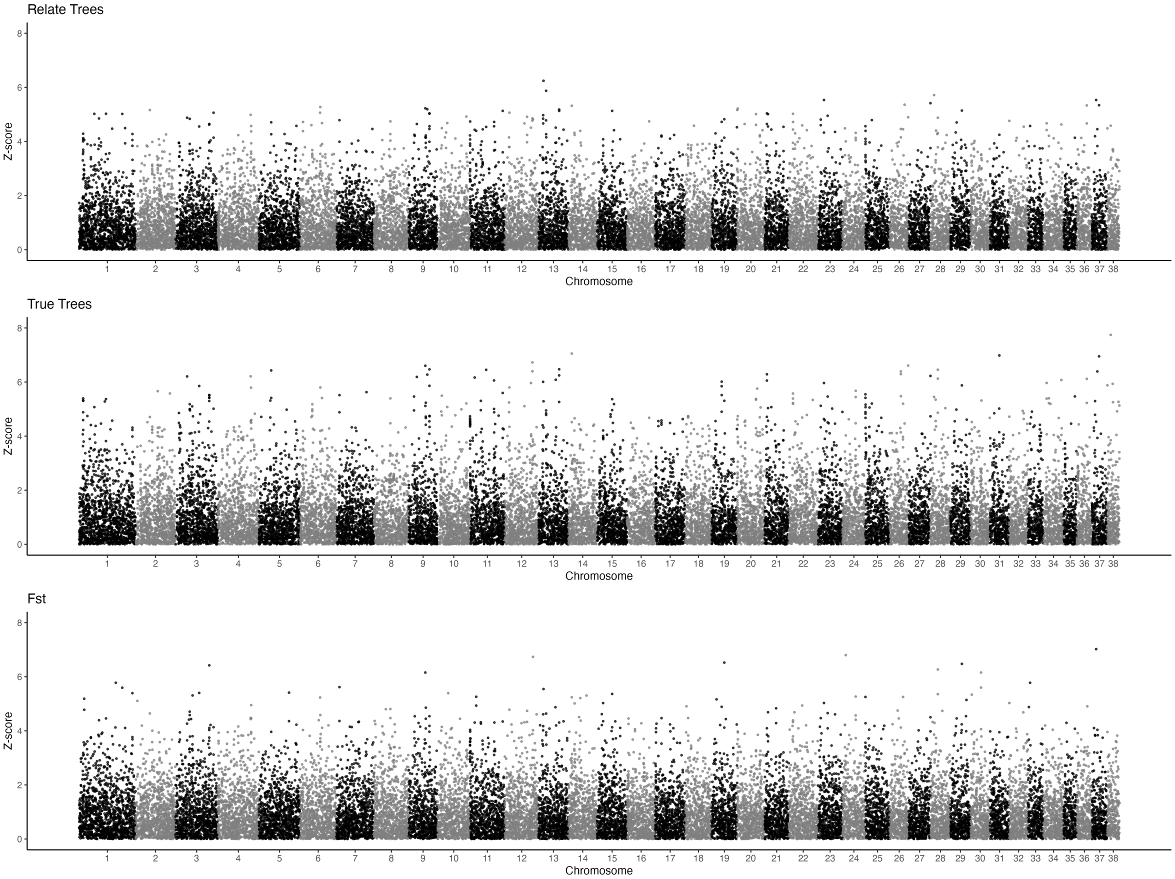


Comparison of TwigScan results for dog and wolf genomes simulated under neutrality, as calculated in 50kbp windows. Shown for z-score > 0 only.

#### SI Figure 12 iHS and XP-EHH scans

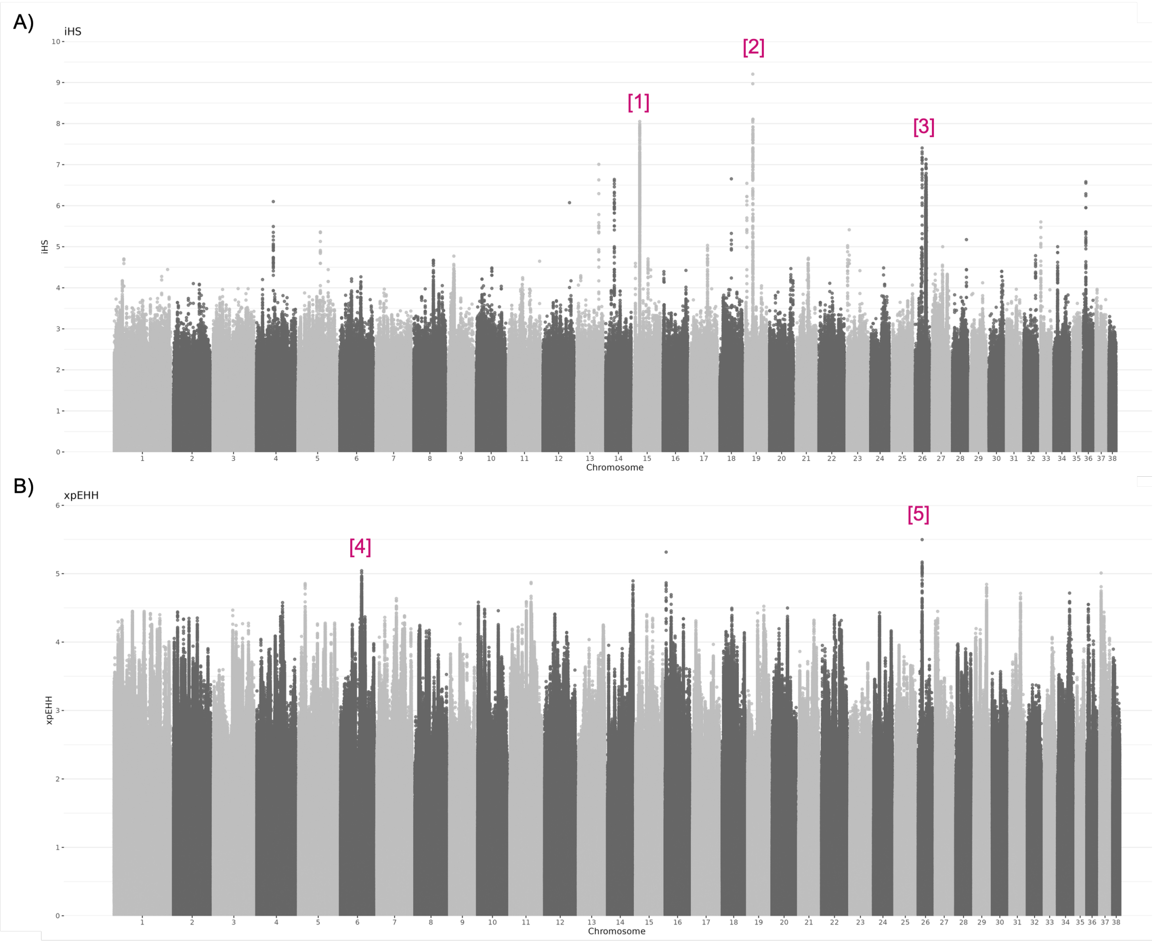


**a,** Normalised absolute iHS as calculated across dogs in the 722g dataset. **b,** Normalised absolute XP-EHH as calculated between dogs and wolves in the 722g dataset. Both were pruned to remove SNPs within 5Mb of the start and end of the chromosome. We labelled notable peaks: [1] indicates a hit identified in this iHS analysis only, overlapping with olfactory receptors annotated in the human genome. Peaks [2] and [3] indicate the chr19 and chr26 hits identified in the Relate analysis, respectively. The chr6 peak [4] is the *RNPC3* hit identified in the TwigScan analysis. The [5] peak indicates a separate hit on chr26 (3.5Mb upstream to the chr26 hit identified by Relate), which does not overlap any known annotated genes.

#### SI Figure 13  Cross-species relationships at RNPC3 and EPAS1

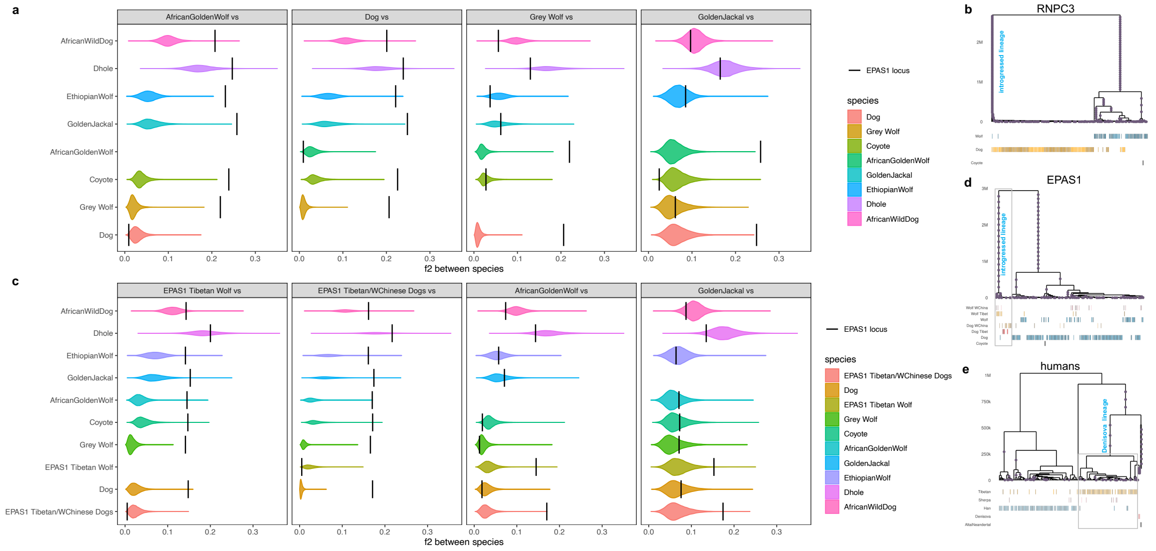


**a,** Distribution of pairwise $f_2$-statistics in 50kb windows, between African golden wolves, dogs, golden jackals, and grey wolves to other wolf-like canids (Supplementary Information). The 100kb window containing the *RNPC3* gene is shown as the black line, indicating that dogs and African golden wolves are diverged at this locus, while golden jackals and grey wolves are not unusually related to other canids. **b,** Relate genealogy at the *RNPC3* gene (47Mbp), corresponding to the highest $f_2$ peak in **a**. **c,** Same analysis as in a, but instead for the *EPAS1* locus which is shown as the black bar. We separated Tibetan and Western Chinese dogs and wolves carrying the *EPAS1* adaptive haplotype from other global dogs and wolves. **d,** Genealogy at the *EPAS1* gene (48,593,806bp, corresponding to midpoint of the gene) in canids from the 722g project. We highlight Tibetan and Western Chinese wolves and dogs who appear to be derived from an introgressed lineage. **e,** Genealogy at the *EPAS1* gene (46,569,191bp) in humans, showing recent Denisovan introgression in Tibetans. Data from Ref. (Lu et al. 2016) and Refs. (Prüfer et al. 2014; Meyer et al. 2012).

#### SI Figure 14 AMY2B copy number variation

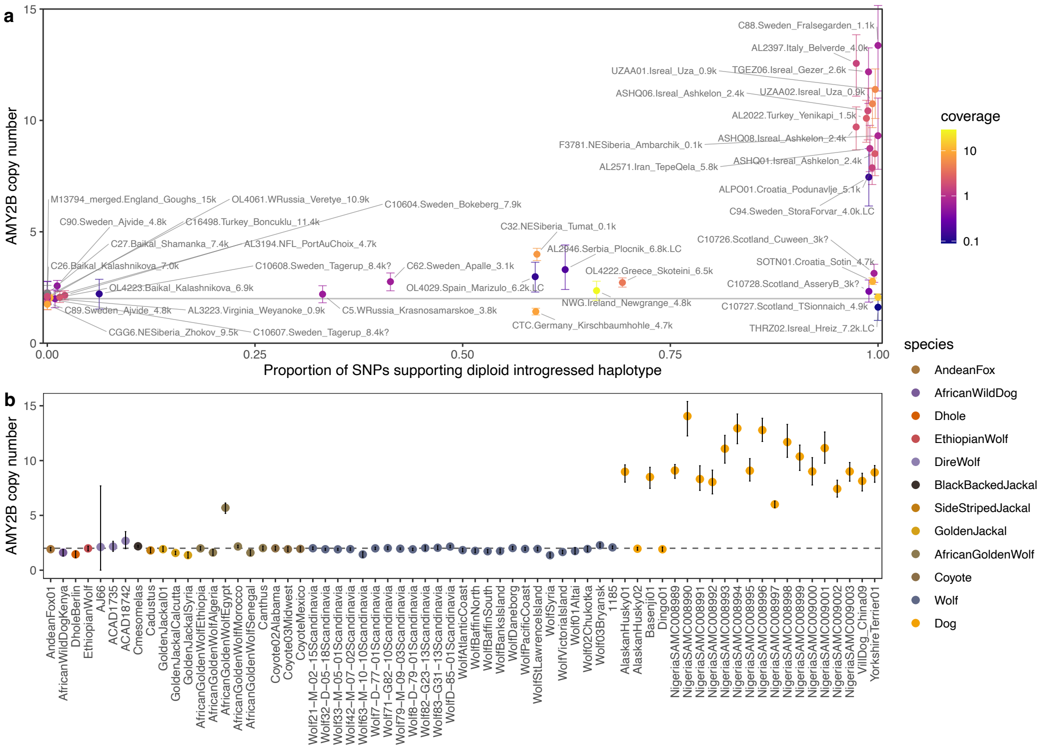


**a,** Copy numbers of *AMY2B* in ancient dogs inferred by aligning to a high-quality grey wolf assembly and then computing relative coverage (Methods), plotted against haplotype carrier status at *RNPC3*. We use the proportion of SNPs in perfect LD tagging the introgressed haplotype. Colour corresponds to genome-wide coverage. **b,** *AMY2B* copy numbers estimated for modern canids. Error bars show 95% confidence intervals obtained from a block bootstrap with block size 500bp.

**SI Figure 15 Simulation evaluating the Relate selection test over epochs.**

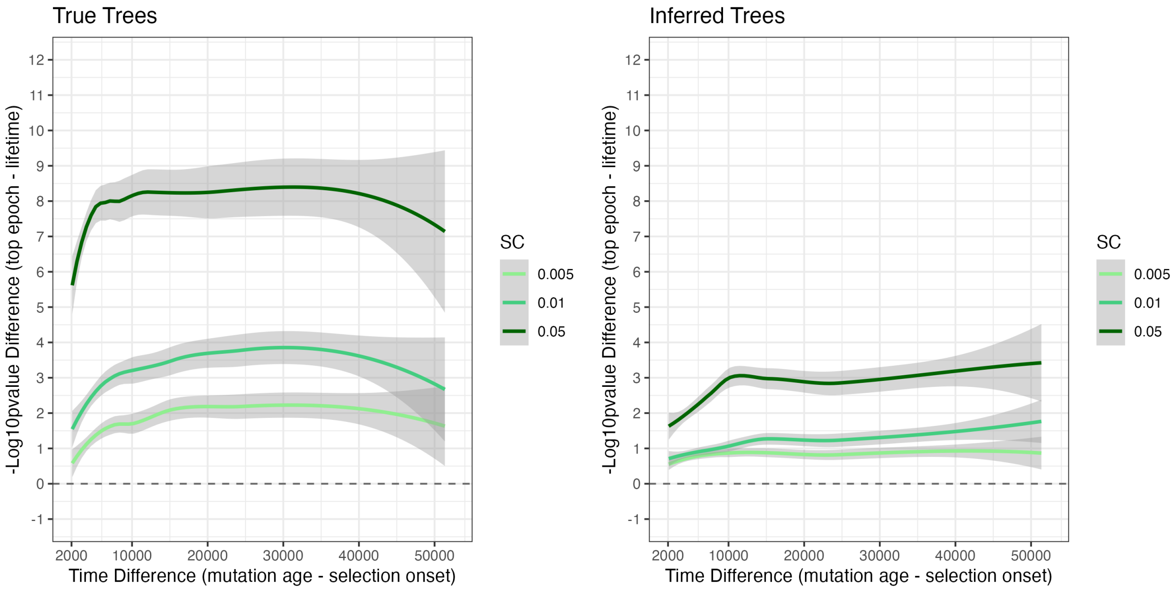


Evidence for positive selection by mutation age on true (left) and Relate-inferred (right) trees in a simulation with variable onset of selection (Supplementary Information). Y-axis shows the difference in $\log_{10}(p)$ between the Relate selection test computed over the lifetime of the mutation and the epoch with the strongest evidence. A positive value indicates increased evidence for selection at the epoch with the strongest evidence. The x-axis shows the difference in generations between mutation age and selection onset in the simulation.

**SI Figure 16 Relate selection test applied to dogs**

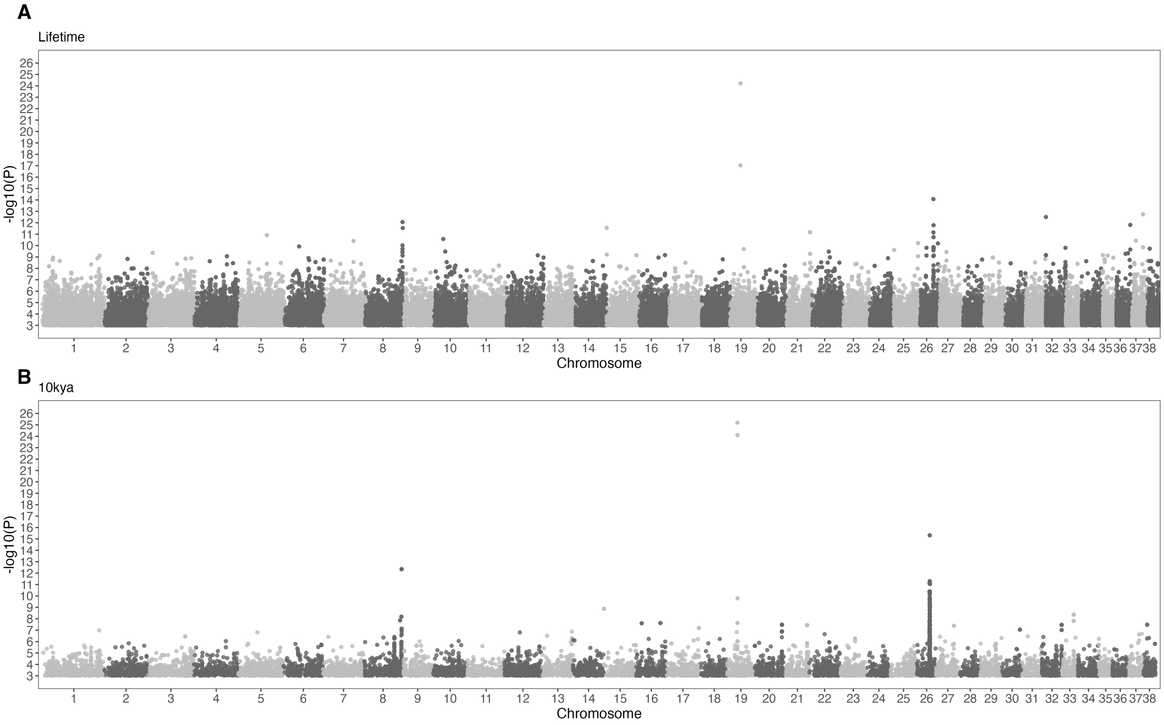


**a,** Manhattan plot of the evidence of positive selection in dogs (722g) calculated using the Relate selection test over the lifetime of the mutation. **b,** Manhattan plot of the evidence of positive selection in dogs (722g) calculated using the Relate selection test over epochs, where we condition on the last 10ky.

**SI Figure 17 Selection statistics and genealogy for the chr26 peak.**

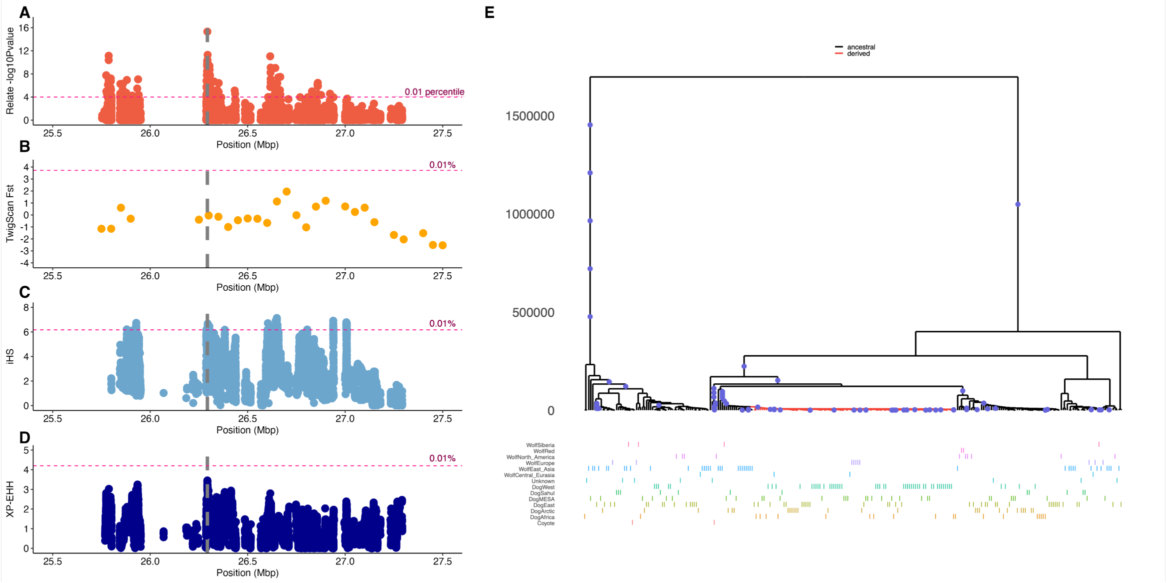


**a,** Relate selection test -$\log_{10}$(p) as calculated from 10kya to present in dogs. **b,** TwigScan $F_{ST}$ between dogs and wolves. **c,** Normalised iHS values for dogs. **d,** Normalised XP-EHH values for dogs and wolves. Dashed grey lines indicate the SNP with the highest evidence in the chr26 peak according to Relate from 10kya to present. Dotted pink lines indicate values at the 0.01% tail for each statistic. **e,** Inferred genealogy of chr26: 26293281 (CanFam3.1; dashed pink line in a-d) in dogs and wolves. Red lineages carry the derived mutation.

**SI Figure 18 Selection statistics and genealogy for the chr19 peak.**

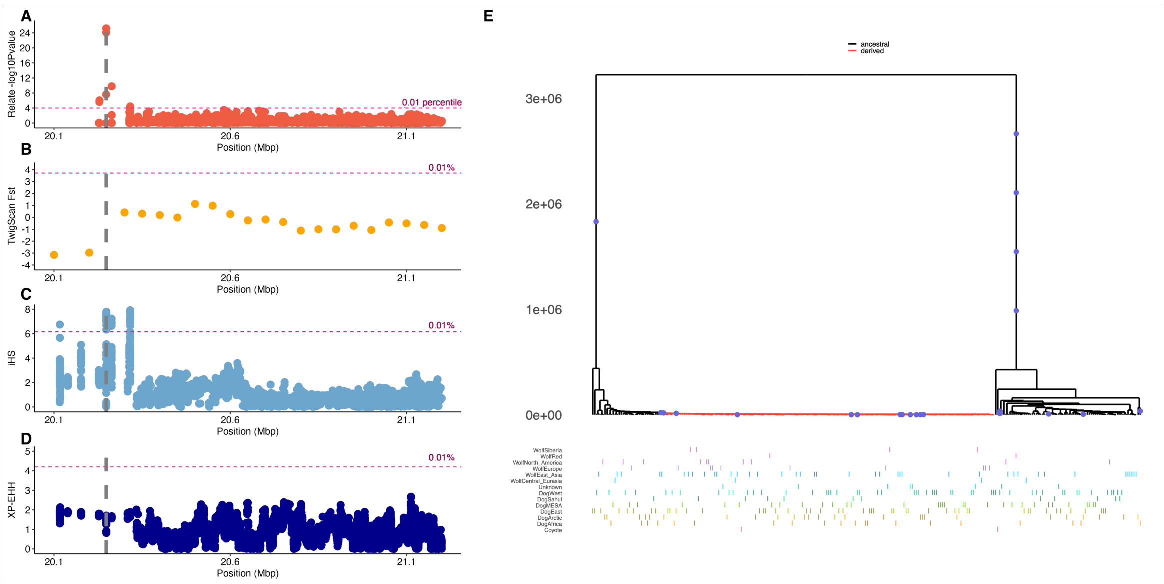


**a,** Relate selection test -$\log_{10}$(p) as calculated from 10kya to present in dogs. **b,** TwigScan $F_{ST}$ between dogs and wolves. **c,** Normalised iHS values for dogs. **d,** Normalised XP-EHH values for dogs and wolves. Dashed grey lines indicate the SNP with the highest evidence in the chr19 peak according to Relate from 10kya to present. Dotted pink lines indicate values at the 0.01% tail. **e,** Inferred genealogy of chr19: 20248039 (CanFam3.1; dashed pink line in a-d) in dogs and wolves. Red lineages carry the derived mutation.

**SI Figure 19 Selection evidence in dogs by frequency in wolves**

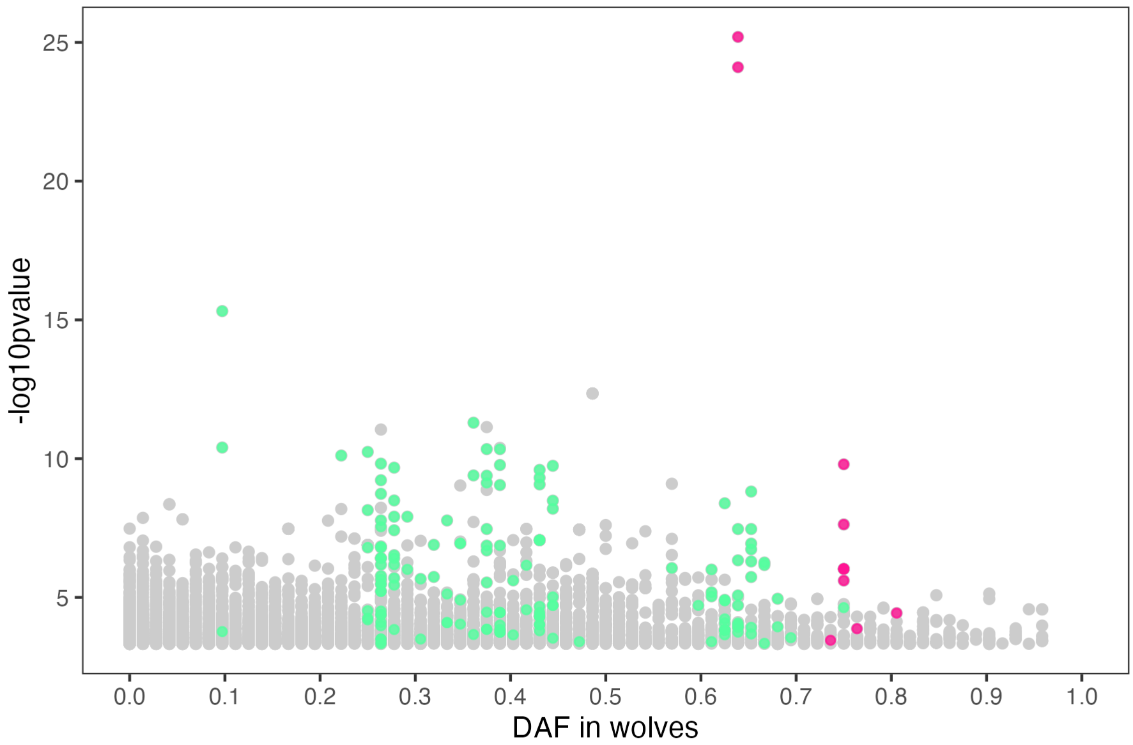


Relate -$\log_{10}$(p) as calculated from 10kya to present plotted against the derived allele frequencies (DAF) of these SNPs in grey wolves (722g dataset). SNPs 20kbp up- and down-stream of the chr19 (pink) and chr26 (green) hits are highlighted.
